# Hydrodynamic Radii of Intrinsically Disordered Proteins: Fast Prediction by Minimum Dissipation Approximation and Experimental Validation

**DOI:** 10.1101/2024.02.05.578612

**Authors:** Radost Waszkiewicz, Agnieszka Michaś, Michał K. Białobrzewski, Barbara P. Klepka, Maja K. Cieplak-Rotowska, Zuzanna Staszałek, Bogdan Cichocki, Maciej Lisicki, Piotr Szymczak, Anna Niedzwiecka

## Abstract

The diffusion coefficients of globular and fully unfolded proteins can be predicted with high accuracy solely from their mass or chain length. However, this approach fails for intrinsically disordered proteins (IDPs) containing structural domains. We propose a rapid predictive methodology for estimating the diffusion coefficients of IDPs. The methodology uses accelerated conformational sampling based on self-avoiding random walks and includes hydrodynamic interactions between coarse-grained protein subunits, modeled using the generalized Rotne-Prager-Yamakawa approximation. To estimate the hydrodynamic radius, we rely on the minimum dissipation approximation recently introduced by Cichocki *et al*. Using a large set of experimentally measured hydrodynamic radii of IDPs over a wide range of chain lengths and domain contributions, we demonstrate that our predictions are more accurate than the Kirkwood approximation and phenomenological approaches. Our technique may prove valuable in predicting the hydrodynamic properties of both fully unstructured and multidomain disordered proteins.

**TOC Graphic:** 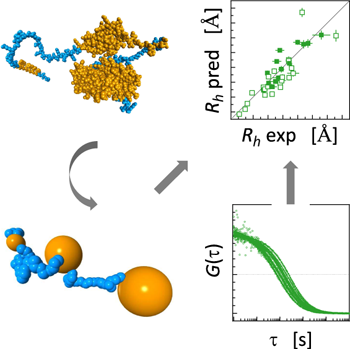

---

Intrinsically disordered proteins (IDPs) constitute an extensive class of biological macromolecules, and their role in the homeostasis of a living cell has been increasingly recognized in recent decades. ^1,2^ The frequency of long intrinsically disordered regions (IDRs) in proteins differs significantly between the kingdoms of life, ranging from 2 % in archaea to 33 % in eukaryotes.^3^ The IDP molecules display different degrees of structural disorder. Their chains can encompass either several folded globular domains, or supersecondary structures connected by flexible linkers, sparse secondary structural elements, or can be completely natively unstructured. Disordered proteins exhibit a notable characteristic – the absence of a stable, well-defined relative spatial arrangement of their fragments. Instead, their equilibrium properties can be described through a broad set of rapidly inter-converting conformers, posing a challenge for analysis, particularly in the context of long chains.^4^

The average geometric properties of IDPs, including their shape and size, are determined by the equilibrium ensemble of conformational states. This equilibrium state is intricately influenced by environmental conditions, ^5^ such as temperature,^6^ ionic strength^7,8^, osmolality,^9^ crowding^10^, post-translational modifications^11^, and the presence of specific molecular binding partners^12^. The formation of transient or more stable non-covalent complexes introduces another non-trivial dependence of the IDP equilibrium geometry on environmental factors.

Because the shape and availability of the binding sites necessary for the interaction of IDP with ligands, other proteins, and nucleic acids are greatly influenced by the environment, IDPs often act as higher-order regulators in key cellular processes such as gene expression^11,13^ signaling^2,14^, or extracellular biomineralization ^15^. It is the different conformations of these flexible proteins, which enable IDPs to perform their multiple functions^1^. In particular, it is worth emphasizing the important roles of IDPs in health and disease, *e.g.*, the role of the p53 protein as a tumour suppressor^16^, mutations of which are often responsible for human cancers, the function of 4E-BPs in the inhibition of eukaryotic translation initiation^11,17–19^, the significance of GW182 protein in the recruitment of the multi-protein machinery necessary for microRNA-mediated gene silencing^20–22^, or the importance of Tau, FUS, and *α*-synuclein proteins in neurodegenerative diseases^23,24^. Since the elastic properties of these biomolecules are responsible for the proper functioning of IDPs in the cellular context, *i.e.* for the association of complexes and the formation of biomolecular condensates via liquid-liquid phase separation such as, e.g., RNA-processing membraneless organelles^25,26^ much attention is paid to the hydrodynamic properties of IDPs. Experimental techniques, such as analytical ultracentrifugation (AUC), size exclusion chromatography (SEC), pulsed-field gradient nuclear magnetic resonance (PFG–NMR), dynamic light scattering (DLS), and fluorescence correlation spectroscopy (FCS), offer insights into hydrodynamic parameters (as reviewed in^27^). However, due to the distinct limitations of each experimental approach, ongoing research aims to devise phenomenological methods for calculating the hydrodynamic radius (*R_h_*). These methods may involve deriving *R_h_* from the radius of gyration determined by small-angle X-ray scattering (SAXS)^28,29^ or exploiting the conformational backbone propensity of IDPs.^30,31^

Simultaneously, significant effort is being invested in developing numerical models that extract the characteristics of IDPs from conformational ensembles obtained using molecular dynamics (MD) simulations or energy minimization algorithms^32–41^. However, the molecular flexibility of IDPs introduces substantial complexities when determining their hydrodynamic properties. Two main issues here are the large number of degrees of freedom and the long timescales of relaxation of the internal coordinates of the molecules. These factors prohibit direct calculation of the experimentally relevant long-time diffusion coefficient from either molecular or Brownian dynamics trajectories. One popular approximation that circumvents this difficulty is to assume that the macromolecule is rigidly frozen in one of a large number of possible conformations. Transport properties are then calculated by treating the molecule as a rigid body, and the results are averaged over an equilibrium ensemble^42–45^. Nevertheless, the validity and accuracy of this approximation remain uncertain. Additionally, the generation of conformational ensembles can be a bottleneck for long chains (beyond approximately 300 amino acid residues) because it requires time-consuming MD simulations and/or the construction of new databases of short peptide conformations.

There is, therefore, a strong need to develop a numerically efficient solution that would enable reliable calculation of the long-time diffusion coefficient of any long-chain IDP, such as one with 1000 amino acid residues, solely based on its sequence information. In this study, we introduce a new theoretical approach to both generating conformational ensembles of IDPs and calculating their hydrodynamic properties. This method enables a swift estimation of the diffusion coefficient for long IDPs in a matter of minutes, with superior accuracy compared to existing methods. This assertion is substantiated through rigorous testing of the model on a diverse set of experimental results obtained for 43 proteins. The dataset includes both literature data and *R_h_* values measured for a set of new IDP constructs using FCS under mild conditions (see Supporting Information). We present our results in terms of the hydrodynamic radius of a molecule, *R_h_*. This radius represents the size of a solid sphere that possesses the same translational diffusion coefficient, *D*, as the given molecule under identical buffer conditions. Therefore, *R_h_* = *k_B_T/*6*πηD*, where *T* is the temperature and *η* is the viscosity.

An important observation by Fixman^46,47^ is that the diffusion coefficient of a flexible macromolecule is time-dependent, with well-defined short- and long-time limits. The disparity between the two is attributed to the effects associated with relaxation of the internal coordinates of the molecule, as well as rotation of the macromolecule as a whole. ^46,48,49^ The positivity of the dissipation rate in the system implies that the long-time diffusion coefficient (*D_l_*) is always smaller than the short-time diffusivity (*D_s_*).^47^ The focus of theoretical approaches should be the determination of the former quantity, as it is the one measured in experiments utilizing techniques like FCS, AUC, or DLS. Unfortunately, the calculation of *D_l_* is significantly more challenging than that of *D_s_*because it involves the computation of time-dependent quantities, such as the memory function, which describes the relaxation effects. An additional point to keep in mind is that the value of the short-time diffusion coefficient depends on the choice of the point that one tracks.^49–52^ In contrast, the long-time diffusivity is independent of the choice of reference point.^53^

The methods for predicting the diffusion coefficient can be broadly split into three categories: atomistic, phenomenological, and coarse-grained. For small proteins, high-resolution, atomistic MD methods can be used,^54^ but they require either simulating the surrounding water molecules explicitly, which is very computationally intensive, or an implicit solvent scheme. In the case of implicit solvent methods, addressing hydrodynamic interactions between distant parts of the molecule^55–58^ and thermalization^59^ pose significant challenges. Additionally, even for the smallest proteins, it is prohibitively difficult to obtain statistically meaningful data over the 10-100 millisecond scale, which would enable the direct computation of the long-time diffusion coefficient.

The other extreme are phenomenological models that predict *R_h_* from the number of residues *N* and possibly other parameters, such as total charge or amino acid composition. Theoretical considerations of Rouse, who modelled a protein as a Gaussian chain^60^ gave foundation to the power law relationship *R_h_* ∼ *N* ^1*/*2^. The classical Rouse model employs random displacements between the monomers. If we assume complete independence of displacements between each consecutive pair of monomers, the central limit theorem dictates that as *N* approaches infinity, the squared end-to-end distance should conform to a scaled *χ*^2^(3) distribution. Consequently, the dimensions of such an idealized chain are expected to scale with ^√^*N*. Later work of Zimm included the effect of excluded volume^61^, which resulted in the scaling *R_h_* ∼ *N^γ^* with *γ* = 0.588.

Phenomenological size–length relationships that include other variables involve a number of fitting parameters. As a result, their range of applicability outside of the fitting dataset is difficult to assess. An alternative phenomenological approach proposed by Pesce *et al*.^29^ employs the radius of gyration obtained from SAXS experiments to estimate *R_h_*. This is substantiated by the observation that within the Kirkwood-Riseman approximation^62^ *R_h_* and *R_g_*share the same scaling relationship with *N* as long as the pair-displacement distribution converges under appropriate scaling to a Gaussian for large *N*.

Finally, coarse-grained models, like our method, employ larger units (typically one or two per amino acid residue) as building blocks for the structure prediction scheme, along with approximate interaction potentials between subunits, to simulate the equilibrium ensemble of configurations for a given molecule. These configurations are then combined with an approximation of the hydrodynamic properties to compute the diffusion coefficient. Essentially, the computation of the latter for elastic macromolecules addresses two interconnected challenges: predicting the conformations of molecules based on available biochemical data and then using these conformations to predict hydrodynamic properties. The different exponents in the power-law relationships of Rouse^60^ and Zimm^61^ demonstrate that even the most basic method for approximating configurations must take into account excluded volume interactions.

A software that can accommodate excluded volume interactions for a disordered chain is Flexible Meccano (FM).^34^ In addition to volume exclusion, it considers the distribution of Ramachandran angles determined from crystallographic protein structures when sampling conformations. However, FM treats the entire chain as unstructured, so it cannot be used to model proteins that possess both globular and unfolded segments, which are in fact much more common than fully unstructured chains. Unfortunately, FM has a closed license that precludes necessary modifications to accommodate folded regions of proteins.

The complex angle distributions used by FM are crucial when computing NMR parameters that are sensitive to short-range details of the pair-distribution function, such as residual dipolar couplings, paramagnetic relaxation enhancement, or J-coupling. However, upon closer examination, the pair-distance distribution generated by FM and a simpler model presented in this paper, globule-linker model (GLM; described below), become virtually identical for amino acids separated by more than 15 residues along the chain.

The highly localized differences between structures at small sequential distances have a minimal influence on the estimations of *R_h_*. It is important to recall that for amino acid residues separated by a distance *r*, the dipolar coupling decays as *r^−^*^3^, while the decay rate of hydrodynamic interactions (HI) is only *r^−^*^1^. Therefore, HI are long-range and less sensitive to near-neighbor distributions, with contributions to the diffusion coefficient of near-neighbors and far-neighbors being *O*(*N*) and *O*(*N* ^2*−γ*^) = *O*(*N* ^1.4^), respectively.

Guided by these considerations, we have implemented the simplest extension of Zimm’s chain — the globule-linker model (GLM), designed to comprehensively represent IDPs containing globular domains connected by unstructured fragments. In the model (Figure 1 A-C), we represented the protein as an assembly of spheres of different sizes. Disordered segments of length *N* were modeled as chains of *N* identical spheres, each with a diameter equal to the *C_α_*-*C_α_* distance, while structured domains were represented by single spheres, each of them described by their mass (*m*). The size of a sphere representing a structured domain was computed as *R_h_* = (3*m/*4*πρ*_globular_)^1*/*3^ + *a*_hydration_ with *ρ*_globular_ = 0.52 Da / Å^3^,^65^ and the single layer hydration shell taken to be *a*_hydration_ = 3 Å thick. Within this approach, information about domain boundaries in the protein sequence is sufficient to construct an appropriate bead approximation of the IDP. The identification of protein sequence fragments to be treated as ordered regions and mimicked by larger beads in the GLM model was done using Disopred3.^66^ The fragment was assumed to be ordered if the disorder probability *P* was less than 50 % for at least three subsequent amino acid residues, including loops linking such fragments but not exceeding 14 residues.^67^

**Figure 1.**
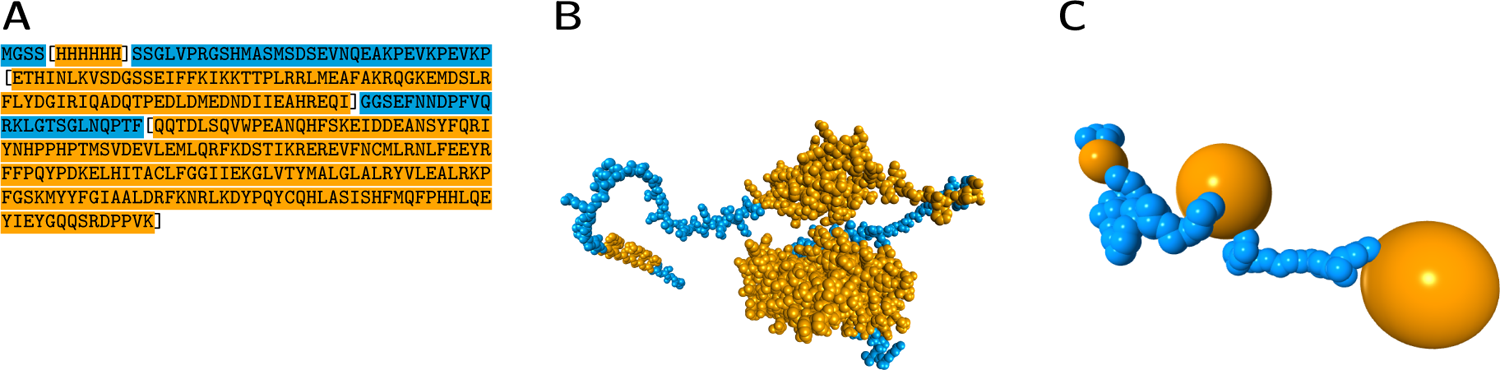
Construction of the coarse-grained globule-linker model (GLM) for an illustratory IDP, H_6_-SUMO-CNOT1(800-999), containing three ordered domains of different sizes (no. 28 in Table S1). **A)** Sequence with highlighted ordered (orange) and disordered (blue) segments, and domain boundaries marked by square brackets. **B)** Representative full atom conformation generated by AlphaFold2 (for visualisation purposes only, ^63,64^ beads with van der Waals radii, hydrogen atoms omitted for clarity); ordered clusters (orange) form dense blobs connected with linkers (blue). **C)** Visualisation of a representative configuration generated using the GLM method where beads are displayed with their hydrodynamic radii.

Using a recursive approach, it is possible to generate GLM conformations with a time complexity of *O*(*N* ^1+*γ*^), which provides a satisfactory ensemble for the largest of the proteins considered here in under a minute using only a personal computer (a single thread at 1.8GHz). The speed of the recursive approach should be contrasted with an iterated one where steps are simply added one by one, and intersecting chains are discarded. This easier-to-implement method is characterized by a time complexity of *O*(exp(*N*)), which becomes prohibitively slow for chains with *N >* 20.

We have transformed the sampled conformations into a hydrodynamic model by increasing bead sizes in the disordered fragments to *R*_disordered_ = 4.2 Å, corresponding to the median value for all aminoacids. ^68^ In the resulting hydrodynamic model of linkers, neighbouring beads show substantial overlaps, requiring a careful treatment of the mobility matrices (see^69^ for details). Note that the value of *R*_disordered_ has only a minor impact on the final results, since the hydrodynamic radius of long slender filaments depends logarithmically on their thickness. ^70–73^

To compute *R_h_* from the estimated ensembles we have implemented two algorithms: the Kirkwood formula, and minimum dissipation approximation (MDA) method of Cichocki *et al*.^53^ Within the first approach,^74^ the hydrodynamic radius of a macromolecule is approximated by

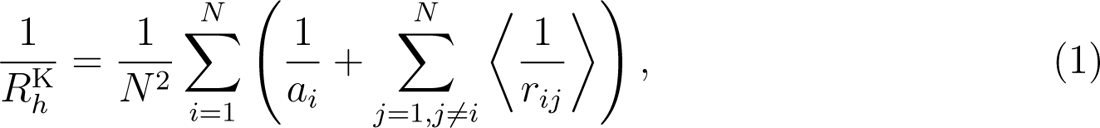

where *N* is the total number of beads in the IDP model, *a_i_* is the hydrodynamic radius of bead *i*, *r_ij_* = |**r***_j_* − **r***_i_*| is the distance between beads *i* and *j*, and ⟨·⟩ denotes the average over the equilibrium ensemble. One can show that this corresponds to the ensemble-averaged short-time diffusion coefficient of the geometric center of the macromolecule, **r***_c_* = *N ^−^*^1^ ^L^*^N^* **r***_i_*. Note that the geometric center fluctuates as the shape of the molecule evolves and does not correspond to any fixed position within it. A simplified form of the Kirkwood formula is often used^39,75,76^

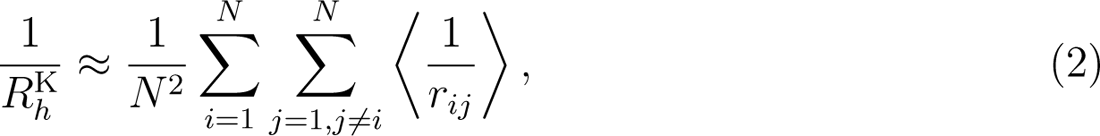

where the single-bead terms 1*/a_i_* are dropped, as their contribution becomes negligible in the large *N* limit. This is the form that we will also use in the present work.

A better estimate of *R_h_*, corresponding to the long-time diffusion coefficient, requires a more in-depth description of the hydrodynamic interactions between the beads. To this end, one introduces the mobility matrix ***µ***,^48^ which links the velocities of the beads with the forces acting on them, according to

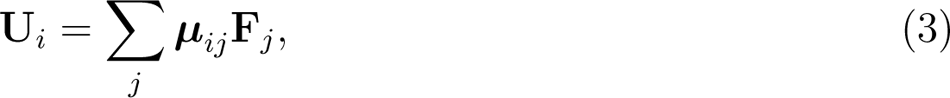

where **U***_i_* is the velocity of bead *i* whereas **F***_j_* is the force with which bead *j* acts on the fluid. Based on the mobility matrix, one defines a matrix **A** indexed by the bead labels (*i*, *j*), *A_ij_*= 2*πη*Tr ***µ****_ij_* and its inverse **B** = **A***^−^*^1^. One can then construct the MDA^53^ for *R_h_* as

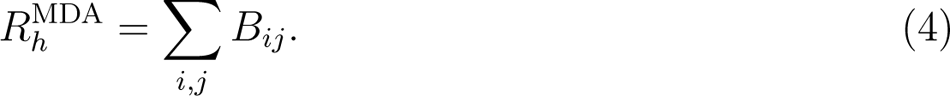

Note that the above formula is general, and can be used for different models of hydrodynamic interactions - both simple models (*e.g.* Oseen or Rotne–Prager far-field approximation^77^) or in more sophisticated approaches, like the multipole expansion method^78,79^. In this work, we use the generalized Rotne-Prager approximation for the calculation of the mobility matrix, as described in^80–82^. This approximation is now also available as a Python package, pygrpy.^83^ For non-overlapping beads, the elements of the matrix **A** have then a particularly simple form: *A_ij_* = ⟨1*/r_ij_*⟩ for *i* = *j*, and *A_ii_* = 1*/a_i_*. The formulas for overlapping beads can be found in the Supporting Information.

The MDA corresponds to the calculation of the short-time diffusion coefficient of the diffusion center of a molecule, ^52^ which is a point inside the molecule where *D_s_* is minimal. The position of the diffusion center is **r***_d_* = Σ*^N^ x_i_***r***_i_*, with the weights given by *x_i_* = Σ *B_ij_/* Σ *B_kj_*. Since *D_s_* is always larger than its long-time counterpart, *D_l_*, MDA provides the best estimation for the long-time diffusion coefficient out of all methods that utilize *D_s_* for this purpose. The MDA turns out to be more robust when dealing with large differences in the sizes of beads used to model constituent parts of the macromolecule, because in such cases the equal weights of the geometric center of the macromolecule differ significantly from the optimal weights of the diffusion center.

We combined each method of generating conformers with each method of computing *R_h_*, which resulted in four different theoretical approaches, the predictions of which (Table S2) were then compared with experimental data. For this purpose, we have obtained 15 new IDP constructs covering a wide range of chain lenghts, folded domain contents and charge states, and determined their *R_h_* using FCS (Figure 2 and S2, S3, S4, S5, S6; for further experimental details see Supporting Information).

**Figure 2.**
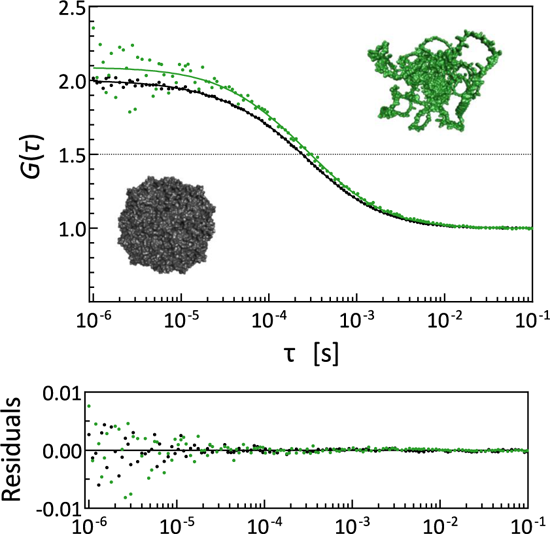
Examples of normalized FCS autocorrelation curves with raw fitting residuals for an intrinsically disordered H_6_-SUMO-GW182SD-mCherry (*N* = 809, *R_h_* = 66 6 Å) (green) in comparison with apoferritin (*N* = 4200, *R_h_* = 58 3 Å) (black). Crystal structure of apoferritin (pdb id. code 2w0o^84^) and putative conformation of H_6_-SUMO-GW182SD-mCherry predicted by AlphaFold ^63^ are shown for illustration purposes, preserving relative sizes of solvent accessible surfaces of atoms.

**Figure 3.**
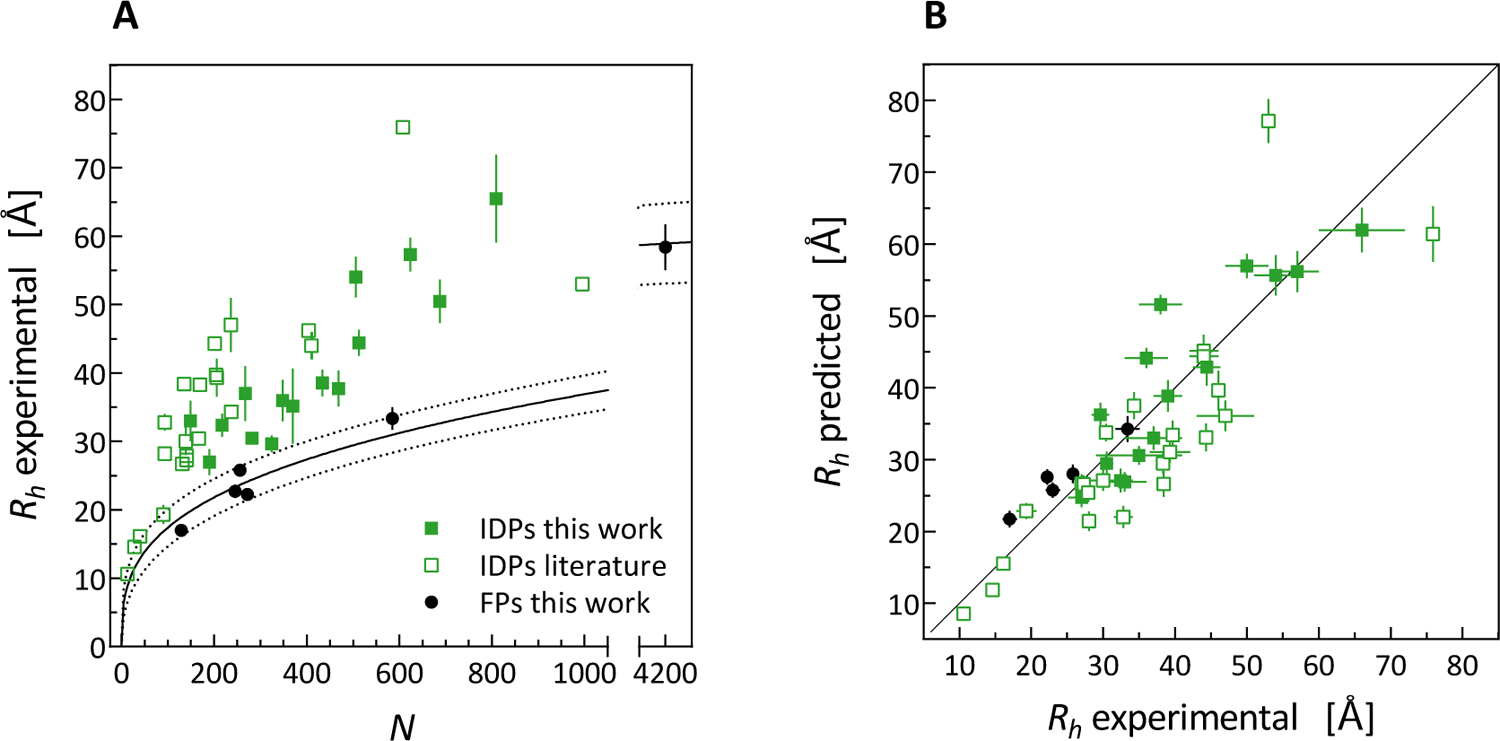
**A)** Experimental *R_h_* values plotted against number of amino acid residues in the protein chain, *N*, and power-law curve fitted to *R_h_* of folded proteins (FP) together with 95% confidence band. **B)** Direct comparison of the predicted vs. measured *R_h_* values for all of the proteins modeled using the MDA+GLM approach.

The experimental benchmark set (Table S1) was thus composed of both the new FCS measurements, and *R_h_* selected from the literature based on the following criteria: the proteins had sequences could be unambiguously identified in the literature or in the UniProtKB database, were measured at well defined, mild conditions (temperature of 20-26 *^◦^*C, buffer of pH 7-8, ionic strength corresponding to 75-300 mM NaCl), and their hydrodynamic radii were determined directly from appropriate experiments without conversions from other experimental quantities, such as *R_g_*.^29,85–102^ This is, to our best knowledge, the largest benchmark set encompassing experimental *R_h_* for 38 IDPs and 6 globular model proteins, measured at comparable conditions.

The results of tests performed for our four theoretical approaches against the bench-mark set are gathered in Table 1, and Figure 4 shows a visual comparison of the deviations between theory and experiment. Additionally, we provide power law fits^104,105^ and power law fits of Marsh *et al*.^103^ for comparison of the prediction accuracy (Table S3).

**Figure 4.**
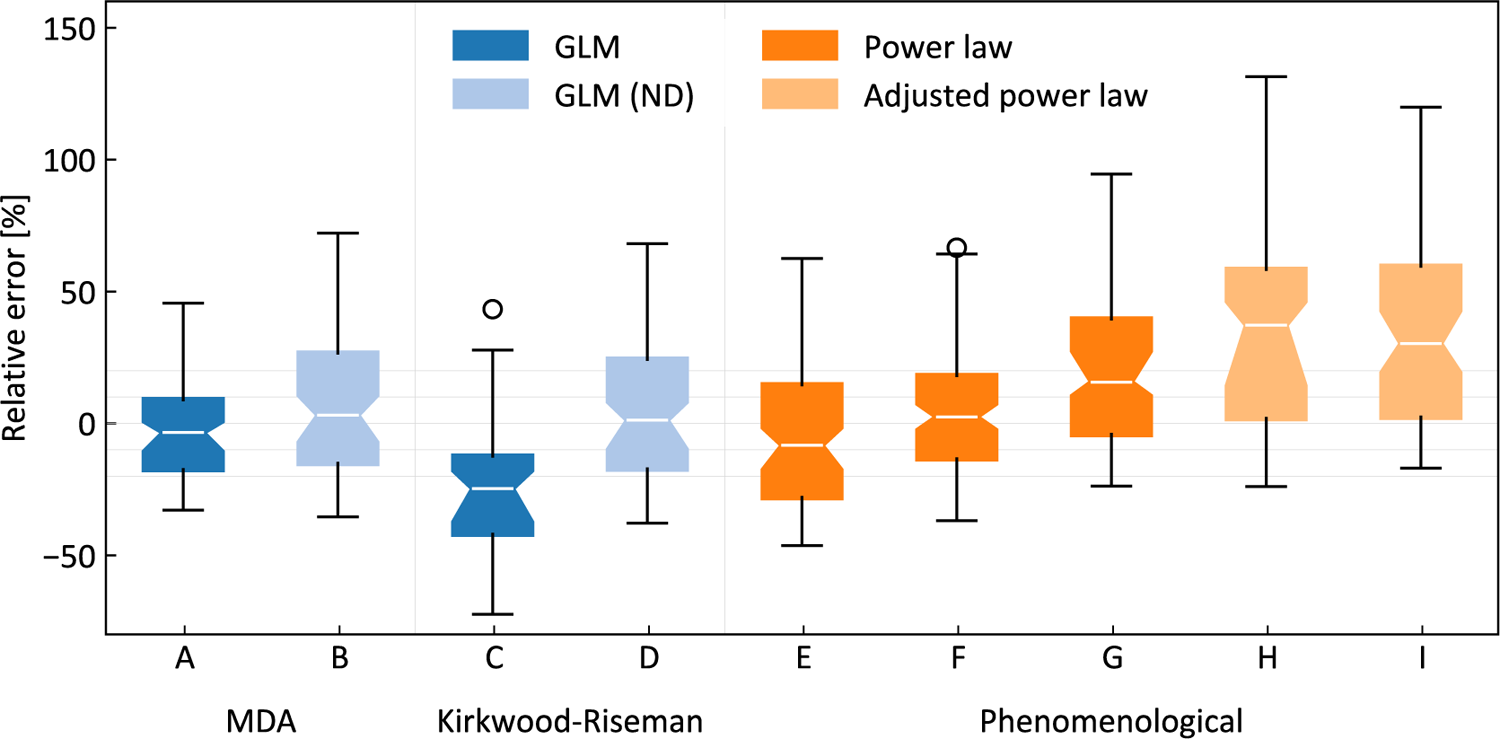
Comparison of different methods of estimation of *R_h_*. Boxes show interquartile range with median confidence bands marked by notches. MDA with GLM ensemble generation (**A**) performs best on the IDP benchmark set with standard errors of 18.15 % and 7.09 Å (compared to 24.80 % and 8.46 Å for a simple power law). Methods based on the Kirkwood-Riseman *R_h_* estimation (**C,D**) typically underestimate hydrodynamic size of the molecule. Power law fits with one free parameter (**E**) and two free parameters (**F**) evaluated using leave-one-out cross validation are compared with the formerly reported power law^103^ (**G**) and a sequence-based model^103^ (**H**) which takes into account total charge of the molecule, and a model based on polyproline II structure propensities^30^ (**I**). Methods with no knowledge about the presence of domains in the IDP (ND; **B,C**) significantly overestimate the hydrodynamic size of the molecule. Domain data can be incorporated into our ensemble generation engine leading to more accurate estimates of *R_h_* (**A**). Note that experimental uncertainty also contributes to the errors presented above and in Table 1.

**Table 1.**
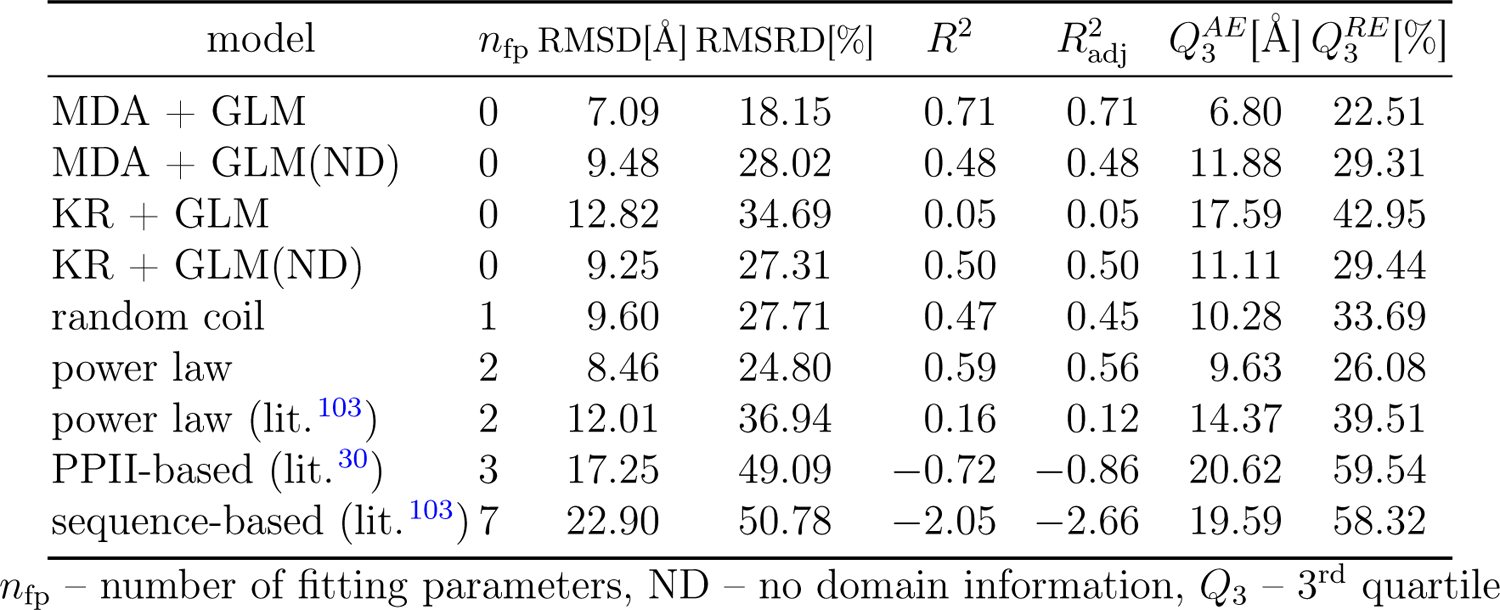
Comparison of error statistics of various models.

We compare the accuracy of the previous and new model under six metrics (Table 1): the square root of mean square deviation (RMSD), square root of the mean square relative deviation (RMSRD), Pearson’s coefficient (*R*^2^), Pearson’s coefficient adjusted for fitting parameters (*R*^2^), 3^rd^ quartile of the absolute error *Q^AE^*, and 3^rd^ quartile of the relative error *Q^RE^*. Whenever a fitting procedure is required, we use leave-one-out cross-validation to compute error metrics. We have chosen to test the relative deviations as well in order to reduce the undue weight given to the new, very long sequences in our dataset. Similarly, outlier-robust metrics of the 3^rd^ quartile were included to reduce the impact of a single sequence misprediction on the final comparisons. In all evaluation metrics, the MDA+GLM approach performs the best. Surprisingly, it is the only model that performs better than the power law baseline in any of the evaluation metrics.

Although explicit intramolecular interactions of the amino acid residues are neglected in MDA+GLM approach, the main cause of discrepancies between the experimental and predicted *R_h_* values (Figure S8) appeared to be the intrinsic properties of individual experimental methods, which suffer from typical errors or limitations and are usually not taken into account when reporting the final experimental results. PGF-NMR measurements are the most unambiguous and accurate, but their effective application is limited to smaller proteins (up to 200-300 amino acid residues long) at high concentrations. It is worth noting that the agreement of values of *R_h_* predicted by MDA+GLM with the PGF-NMR results is excellent (Figure S8 C). FCS is the only method that addresses the self-diffusion of molecules at the limit of low concentrations. Raw FCS measurements can be refined to exclude possible oligomerization or aggregation during the experiment based on the count-rates. However, it is impossible to avoid proteolytic instability of proteins and, consequently, the appearance of impurities with a lower molar mass, which may potentially result in apparently lower values of *R_h_* (Figure S8 B). On the other hand, SEC is the easiest approach to remove lower mass impurities, but it involves diffusion of molecules at higher concentrations through a medium with pores of a specific shape under the influence of pressure. An additional common disadvantage is calibration based on *R_h_*of standard proteins determined in various conditions and the lack of appropriate propagation of the calibration experimental uncertainty. Consequently, SEC measurements can be highly scattered (Figure S8 D). The largest outlier in our analysis concerns *R_h_* determined using SEC for fesselin without providing experimental uncertainty (Id. 43, Tables S1, S2, Figure S7, S8 D). The DLS method is the most prone to overestimating experimental values (Figure S8 E), since the presence of even a small number of aggregates with a larger molar mass generates a huge contribution to the intensity of scattered light. Finally, AUC yields sedimentation coefficients, and their interpretation in terms of exact values of *R_h_* requires some assumptions that are not obvious for IDPs, such as *e.g.* partial specific protein volume.^106^ The second largest outlier in our set is the OMM-64 protein (Id. 39, Tables S1, S2, Figure S7, Figure S8 F) with the *R_h_*value determined using AUC, which is very close to the power function curve for completely denatured proteins.^107^

In conclusion, we have presented a simple, first principles model for the prediction of *R_h_* without any fitting parameters and achieved favourable comparison with a large benchmark set. Moreover, due to the relative simplicity of the model, all of the calculations for a given protein can be performed in about a minute on a typical laptop, which is contrasted with MD-based conformer generation methods that require supercomputers and take many days. Furthermore, to our surprise, the GLM-MDA approach demonstrates satisfactory convergence even with ensemble sizes as small as 40 conformers.

Our benchmark set, in which the previously known IDPs were complemented by a set of newly obtained proteins, constitutes a significant step forward in predicting hydrodynamic properties of IDPs. It includes a higher conformational variety, with a stronger emphasis on multidomain proteins, longer chains, and a much wider range of charge states compared to the reference sets used previously.^30,103^ This diversity allows for more reliable testing of theoretical models.

Further developments of the MDA+GLM model are needed to take into account the dependence of *R_h_* on the environmental conditions^6–8^ and the formation of complexes. However, our results clearly demonstrate that the relatively simple globule–linker model for conformational ensemble construction, in combination with the minimum dissipation approximation, can serve as the starting point for developing further phenomenological corrections. These improvements could incorporate factors such as amino acid sequence composition, residue charge, and counterion binding. When using MDA+GLM, all excluded volume effects should be correctly accounted for, with any further deviations hinting at interesting physical and chemical properties of the molecules.

## Acknowledgement

The work of AM, MKB, BPK, MKC-R, ZS, and AN was supported by the National Science Centre of Poland Sonata-Bis grant no. UMO-2016/22/E/NZ1/00656 to AN. The work of RW and ML was supported by the National Science Centre of Poland Sonata grant no. 2018/31/D/ST3/02408 to ML. The authors thank Prof. Nahum Sonenberg and Dr. Marc Fabian for sharing plasmids for some protein constructs and Dr. Joanna Żuberek and Dr. Mateusz Kogut for helpful discussion. The research was performed in the NanoFun laboratories co-financed by ERDF within the POIG.02.02.00-00-025/09 Program.

## Supporting Information Available

Tables S1 S5 Figures S1-S8 Materials and Methods.pdf containing experimental *R_h_*, Figures S1, S2, S3, S4, S5, S6, S7, S8, and additional methodological details. Tables S2-S4.xlsx containing theoretical *R_h_* from GLM, phenomenological *R_h_*, and protein sequences with marked domains.

The programs referenced in this article can be found on GitHub. For reader convenience, an API and a command-line Python utility, glm_mda_diffusion, have been provided.^108^ This package builds upon previous ones, namely pychastic and sarw_spheres, both of which are also accessible on GitHub.

