## Supplementary material for "Hydrodynamic Radii of Intrinsically Disordered Proteins: Fast Prediction by Minimum Dissipation Approximation and Experimental Validation": Tables S1 S5 Figures S1-S8 Materials and Methods

Additional details, materials, and methods

### Materials and Methods

#### Computational details

Fast convergence of the MDA-GLM algorith for computation of $R_h$ values.

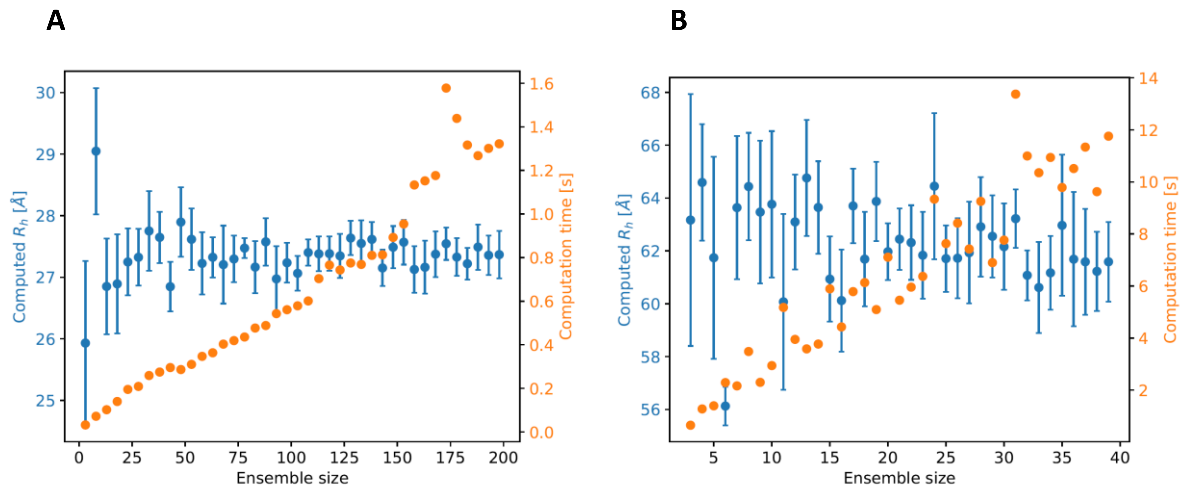


**Figure S1.** Computed $R_h$ value (blue) and computational time (orange) as a function of ensemble size for two cases, A) a small SAP 1A protein (n = 149, id = 13, Table S1) and B) a large $H_6$-SUMO-GW182 SD-mCherry protein (n = 809, id = 42), presented with 2 standard deviations error bars estimated using 10 rounds of bootstrap, included in the computation time. Even for moderate ensemble sizes (N=20), Monte Carlo errors are smaller than hydrodynamic approximation errors.

#### Experimental details

*Chemicals*

The chemicals for protein expression and purification were purchased from Merck (Sigma-Aldrich) and were analytically pure, grade A, or specified for molecular biology. The AF488 NHS ester was purchased from Lumiprobe GmbH. Alexa Fluor 546 NHS ester was purchased from Invitrogen.

*Standard proteins*

Apoferritin, human serum albumin (HSA), α-chymotrypsinogen A, and lysozyme were purchased from Merck (Sigma-Aldrich).

*Protein expression, purification and labelling*

$H_6$-SUMO-CNOT1(800-999), GST-CNOT1(800-999), eIF4E, and eIF4E(28-217) were expressed and purified as described previously[1–5]. The genes for $H_6$-SUMO-SAP 1A, SUMO-mαEGFP-$H_6$, $H_6$-SUMO-GW182 SD ΔRRM, $H_6$-SUMO-AGARP, $H_6$-SUMO-PARN C-mCherry, $H_6$-SUMO-GW182 SD-mCherry protein constructs were ordered from BioCat

GmbH (Heidelberg, Germany). $H_6$-mCherry and $H_6$-mCherry-α-helix were kind gifts from Dr. Joanna Grzyb. The proteins were overexpressed in *Escherichia coli* Rosetta 2(DE3)pLysS and purified in the form of fusion proteins by Ni-NTA affinity. To obtain SAP 1A, mαEGFP-$H_6$, GW182 SD ΔRRM, AGARP, and GW182 SD-mCherry, the fusion proteins were digested from the SUMO-tag by using the His-tagged SUMO Protease (Sigma-Aldrich) or, in the case of mαEGFP-$H_6$, by the untagged CoolCutter SUMO Protease (GeneCopoeia), according to the protocols provided by the manufacturers. The proteins were further purified by anion exchange chromatography by using HiTrap Q or HiTrap SP (Cytiva), depending on the protein construct pI values, followed by size exclusion chromatography (SEC) with use of the Superdex 200 Increase 10/300 GL column (Cytiva) at ÄKTA pure FPLC system (GE Healthcare). The protein purity was checked by the SDS PAGE and depended on the protein construct, ranging from 80% for the coral acid-rich proteins and $H_6$-SUMO-GW182 SD-mCherry, to 99% for eIF4E and model folded proteins,). The identity of all new proteins has been confirmed by mass spectrometry. The sequences of the proteins are given below.

Proteins were labelled by using the AF488 NHS ester according to the manufacturer's protocol (Lumiprobe GmbH ) and purified from the excess of the unreacted dye by Zeba spin columns (Thermo Scientific), multi-step dialysis with use of Pur-A-Lyzers (Sigma-Aldrich), or by another SEC run on Superdex 200 Increase 10/300 GL(Cytiva), depending on the protein properties. The residual presence of the unreacted dye was taken into account in the FCS data analysis as a second component.

*Fluorescence correlation spectroscopy measurements*

The FCS experiments were performed essentially as described previously [6], at Zeiss LSM 780 with ConfoCor 3, in 50 mM Tris/HCl buffer pH 8.0 (at 25 °C), 150 mM NaCl, 0.5 mM EDTA, and 1 mM TCEP or DTT, in droplets of 25-30 μl. The buffer and the samples were filtered through the membrane of 0.22 μm pore sizes immediately before the experiment. The protein concentrations were in the range of 10-20 nM after the filtration. The temperature inside the droplet, $25 \pm 0.5$ °C, was checked after the FCS measurements by means of a certified calibrated micro-thermocouple. A single measurement time was 3 to 6 s, repeated 10 to 100 times in a set. The set of measurements was repeated 3 times in 5 independent droplets.

The structural parameter ($s$) was determined every time with use of AF488 ($D_{AF488}$ = 435 μm$^2$ s$^{-1}$) or Alexa Fluor 546 ($D$ = 341 μm$^2$ s$^{-1}$) in pure water [7], individually for each microscopic slide previously passivated with BSA in the working buffer. The actual solution viscosity was taken into account by comparison of the diffusion time for AF488 or Alexa Fluor 546 in pure water and in the buffer at the same equipment calibration.

The experiments for proteins labelled by AF488, SUMO-mαEGFP-$H_6$, and mαEGFP-$H_6$ were performed at the 488 nm excitation wavelength with a relative Argon multiline laser power of 3 %, MBS 488 nm, BP 495-555 nm. For the mCherry-fused proteins and Alexa Fluor 546 calibration, the excitation wavelength was 561 nm at 2 % relative DPSS laser power, MBS 488/561 nm, LP 580 nm. A dampening factor of 10 % and a dust filter of 10 % were applied.

Photophysical processes of AF488 and fluorescent proteins, mCherry and mαEGFP, were investigated in independent sets of experiments. A relative laser power ranging from 3 to 20 % at 488 nm was used for the AF488 triplet state lifetime measurements. The average lifetime was determined to be about 4 μs. In the case of mCherry and mαEGFP, the measurements were performed in 30 % glycerol to slow down the protein diffusion and extract the blinking [8]. The fraction of mCherry population that undergoes blinking was found to be about 24-28 %

both for the fluorescent protein alone and in the fusion constructs, and about 15 % for mαEGFP.

*FCS data analysis*

The FCS data were analysed by using the Zen2010 software (Zeiss). The raw measurements were closely inspected and refined to exclude possible oligomerization or aggregation of the protein sample in the confocal volume during the experiment. Global fitting of the autocorrelation curve was performed to data sets containing 10 to 50 single measurements. The autocorrelation function for 3D diffusion, including photophysical processes (triplet state for chemical dyes or blinking for fluorescent proteins) was fitted according to the equations [9]:

$$G(\tau) = G_T(\tau) \cdot G_D(\tau) \qquad \text{(eq. 3)}$$

$$G_T(\tau) = (1 + \frac{P_T}{1 - P_T} e^{-\frac{\tau}{\tau_T}}) \qquad \text{(eq. 4)}$$

$$G_D(\tau) = \sum_{i=1}^{n} \frac{\Phi_\mathrm{i}}{\left(1 + \left(\frac{\tau}{\tau_{\mathrm{d,i}}}\right)\right) \cdot \left(1 + \left(\frac{\tau}{\tau_{\mathrm{d,i}}}\right) \cdot \frac{1}{\mathrm{s}^2}\right)^{1/2}} \qquad \text{(eq. 5)}$$

$$\sum_i \Phi_i = 1 \qquad \text{(eq. 6)}$$

where: $G(\tau)$ is the fitted autocorrelation function; $G_T(\tau)$, normalized autocorrelation function for photophysical processes; $G_D(\tau)$, normalized autocorrelation function for the diffusion of $n$ components; $P_T$, triplet state or blinking fraction; $\tau_T$, lifetime of the photophysical process; $\tau_{d,i}$, diffusion time for the $i$-th component; $s$, structural parameter of the confocal volume; $\Phi_i$, fraction of the $i$-th diffusing component.

A one-component model ($n = 1$) providing for the fluorescent protein blinking was fitted for the fusion proteins, and a two-component model ($n = 2$), taking into account the AF488 triplet state and the presence of a residual freely diffusing dye, was used for the chemically labelled proteins. The mCherry and mαEGFP blinking fraction, as well as the AF488 triplet state lifetime determined from the independent experiments were fixed during the global analysis.

The $R_h$ values were determined from the diffusion times, $\tau_d$, providing for the actual buffer viscosity, as follows:

$$R_h = \frac{kT \cdot \tau_d}{6\pi \eta_0 \cdot D_{dye} \cdot \tau_{dye\_buf}} \qquad \text{(eq. 7)}$$

where $\eta_0$ is the viscosity of pure water [10] at the temperature $T$ and $\tau_{dye\_buf}$ and $D_{dye}$ is the diffusion time of AF488 or Alexa Fluor 546 in the buffer at the same calibration.

The numerical regressions were performed by Prism 6 (GraphPad Software).

The total experimental uncertainty was determined according to the propagation rules for small errors[11], taking into account both numerical uncertainty of the fitting, statistical dispersion of the results, and uncertainties of other experimental values used for calculation of the results.

A power function of the number of the polymer units (N) was fitted to the experimental $R_h$ values of folded proteins, determined by FCS (Table S1) according to the equation:

$$R_h(N) = R_0 N^{\nu} \qquad \text{(eq. 8)}$$

The critical exponent value, $\nu$, was calculated as 0.33 ± 0.02, in agreement with the value of 1/3 for a polymer chain packed into a spherical shape, and the $R_0$ was determined as 3.9 ± 0.6 Å, which corresponds to an average $R_h$ value for free amino acids, 3.2 ± 0.4 Å [12].

#### Bioinformatics

Example conformations of the IDPs were generated by AlphaFold 2.0 notebook [13,14]. Protein structures were drawn by using Discovery Studio v3.5 (Accelrys Software).

Identification of the protein sequence fragments to be treated as ordered regions and mimicked by larger balls in the globule-linker model (GLM) was done by using Disopred3[15]. The fragment was assumed to be ordered if the disorder probability P was less than 50 % for at least three subsequent amino acid residues, including loops linking such fragments not exceeding 14 residues[16].

#### Selection of $R_h$ from literature data

The experimental benchmark set was complemented by the $R_h$ values selected from literature. The selected proteins had sequences that could be unambiguously identified in the literature or in the UniProtKB database, were measured at well defined, comparable, mild conditions (temperature of 20 - 26 °C, buffer of pH 7 - 8, ionic strength corresponding to 75 - 300 mM NaCl), and their hydrodynamic radii were determined directly from appropriate experiments without conversions from other experimental quantities, such as $R_g$[17–35].

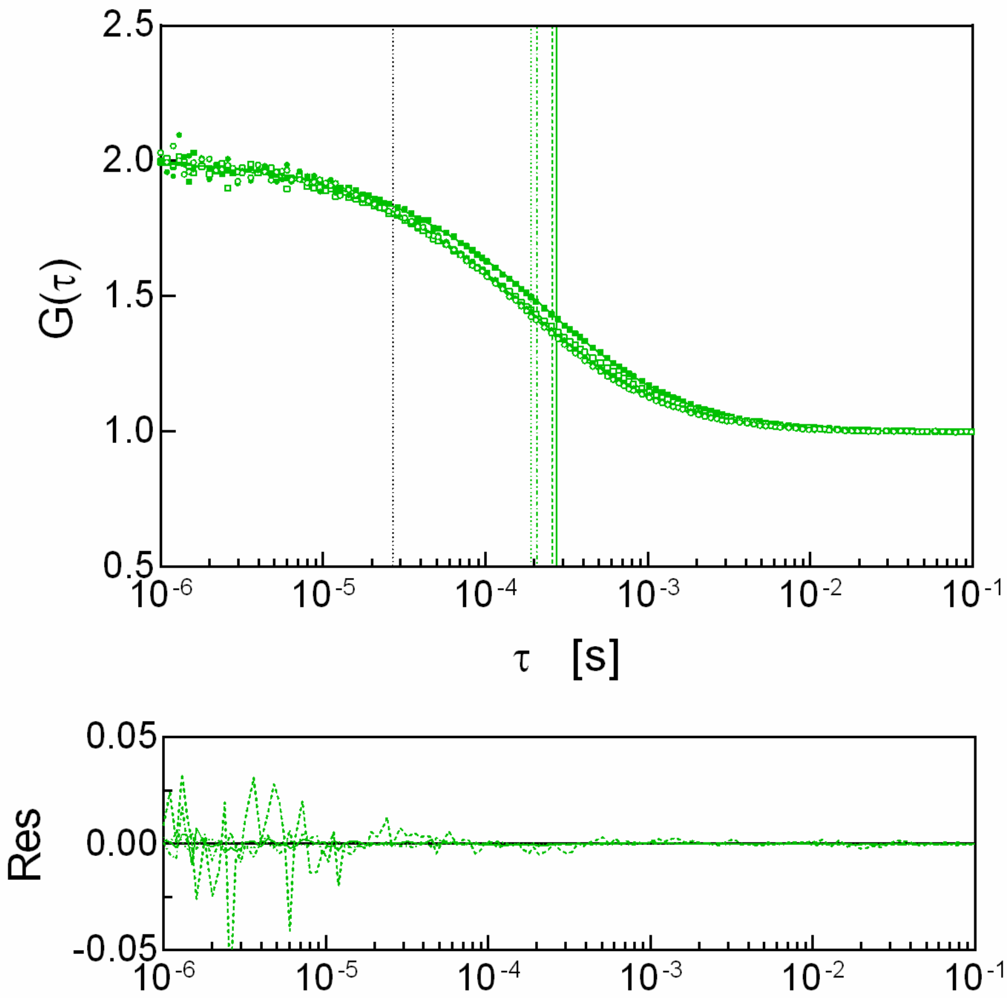


**Figure S2.** Example normalized FCS data and autocorrelation curves for SAP 1A (149 residues) (○), $H_6$-SUMO-SAP 1A (267 res.) (●), AGARP (506 res.) (□), and $H_6$-SUMO-AGARP (624 res.) (■) with their fitting residuals (·····,-··-·-, -----, —, respectively). Vertical lines in the upper panel indicate the diffusion times for the residual free AF488 dye (black ····· line at ~27 μs) and the proteins (green ·····, -··-·-, -----, — lines at 191 μs, 208 μs, 259 μs, and 274 μs, respectively).

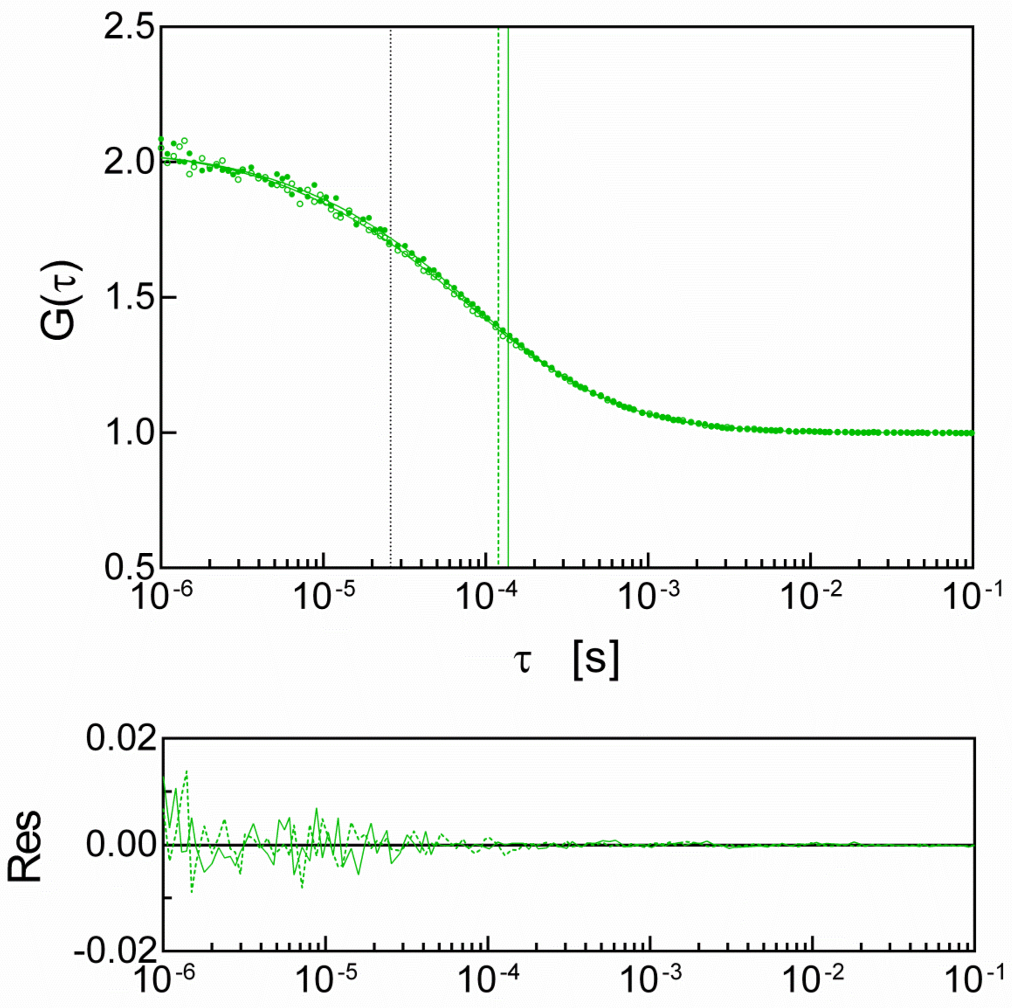


**Figure S3.** Example normalized FCS data and autocorrelation curves for eIF4E(28-217) (○) and eIF4E(1-217) (•) with their fitting residuals (-----, —, respectively). Vertical lines in the upper panel indicate the diffusion times for the residual free AF488 dye (black ····· line at ~26 μs) and the proteins (green -----, — lines at 120 μs, and 138 μs, respectively).

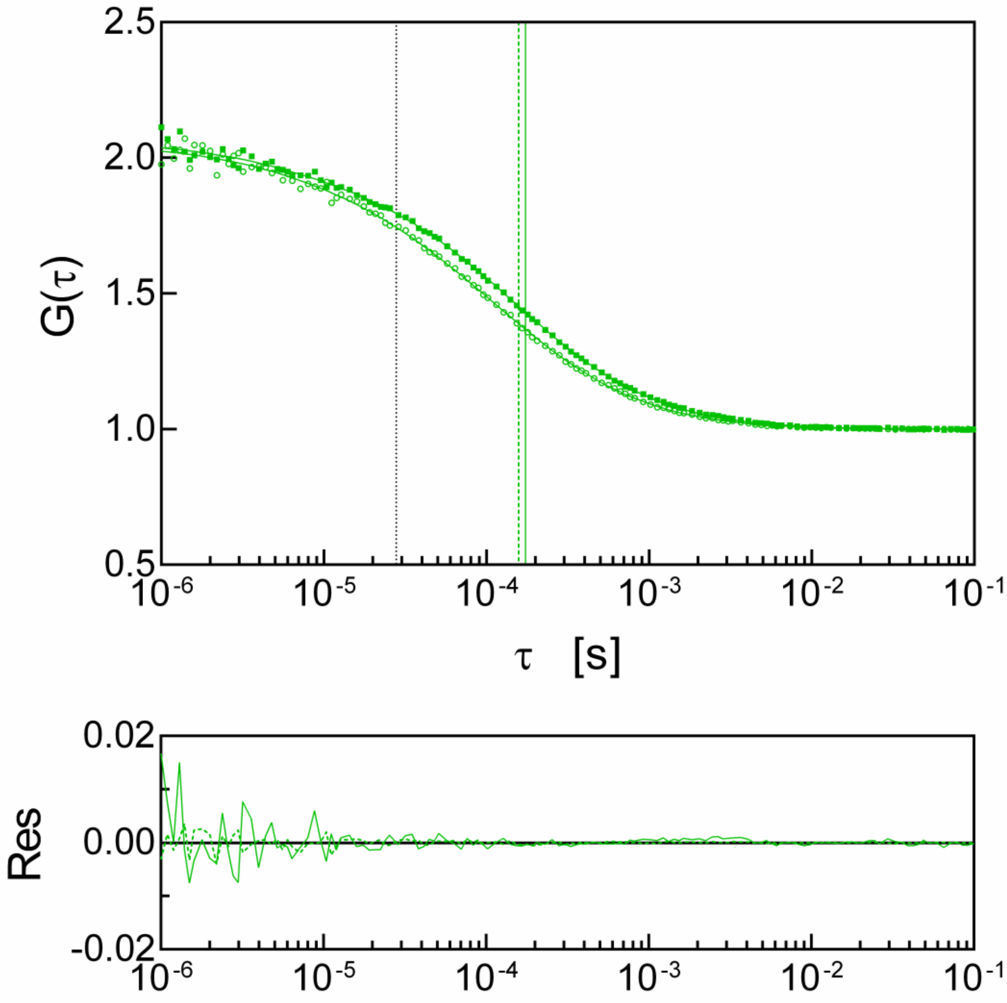


**Figure S4.** Example normalized FCS data and autocorrelation curves for $H_6$-SUMO-CNOT1(800-999) (324 res.) (∘) and GST-CNOT1(800-999) (434 res.) (▪) with their fitting residuals (-----, —, respectively). Vertical lines in the upper panel indicate the diffusion times for the residual free AF488 dye (black ····· line at ~28 μs) and the proteins (green -----, — lines at 158 μs, and 174 μs, respectively).

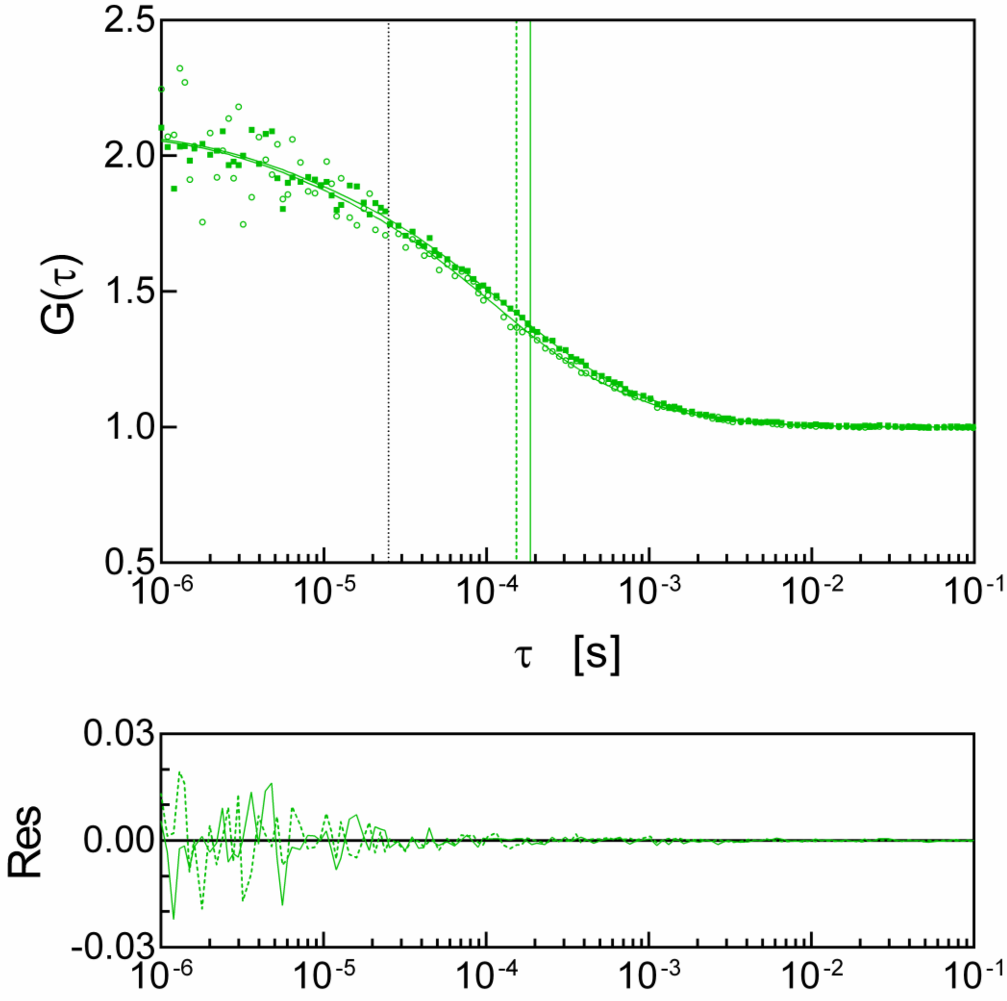


**Figure S5.** Example normalized FCS data and autocorrelation curves for GW182 SD ΔRRM (348 res.) (◦) and $H_6$-SUMO-GW182 SD ΔRRM (469 res.) (▪) with their fitting residuals (-----, ––, respectively). Vertical lines in the upper panel indicate the diffusion times for the residual free AF488 dye (black ····· line at ~25 μs) and the proteins (green -----, –– lines at 153 μs, and 186 μs, respectively).

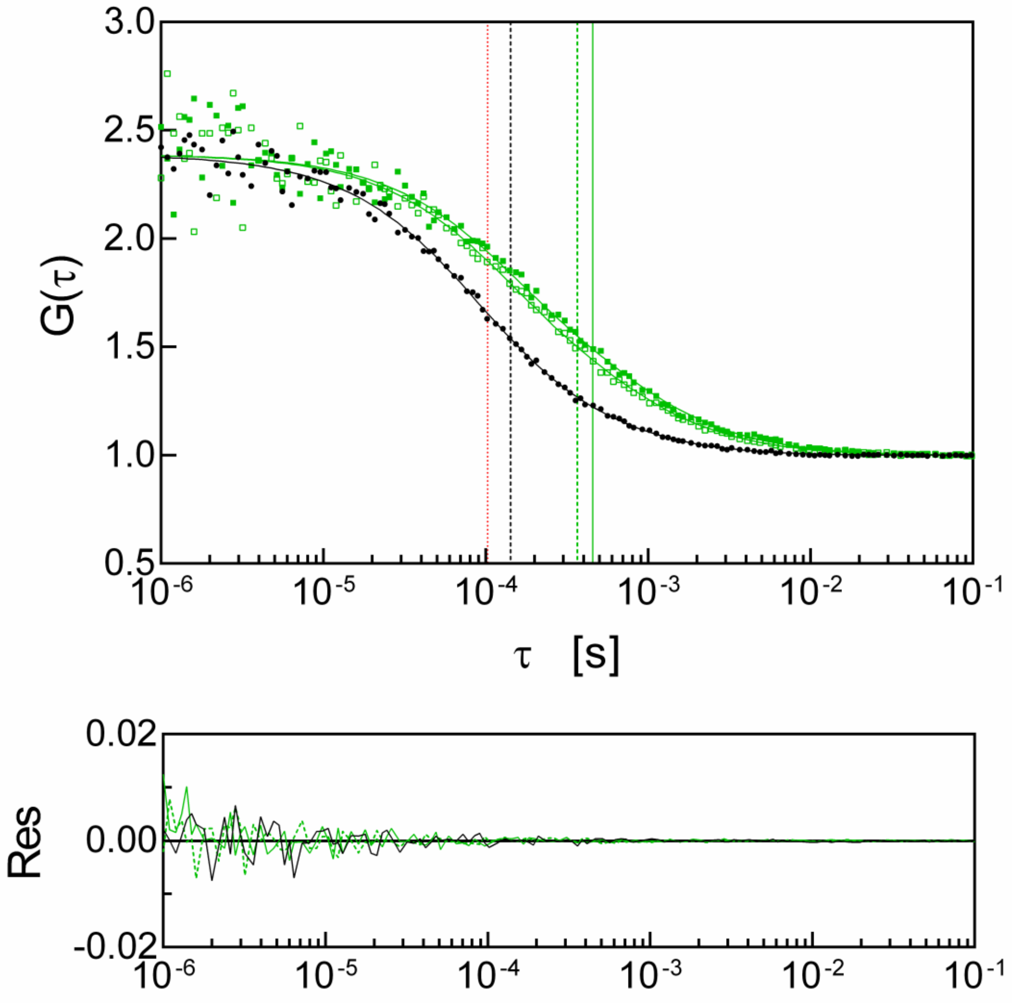


**Figure S6.** Example normalized FCS data and autocorrelation curves for $H_6$-mCherry (256 res.) (black •), GW182 SD-mCherry (688 res.) (809 res.) (green ▫), and $H_6$-SUMO-GW182 SD-mCherry (green ▪) with their fitting residuals (black —, green -----, green —, respectively). Vertical lines in the upper panel indicate the blinking time for the mCherry fluorophore (red ····· line at ~103 μs, 28 % blinking fraction) and the diffusion times for the proteins (black -----, green -----, green — lines at 143 μs, 367 μs, and 458 μs, respectively).

**Table S1.**

Experimental values of hydrodynamic radii, $R_h$, for the benchmark proteins. Most of them are intrinsically disordered proteins (otherwise noticed in the Remarks column). N, number of amino acid residues in the protein chain.

| Id. | Protein name | N | $R_h$ exp [Å] | $\Delta R_h$ exp [Å] | T [°C] | Conditions/ Remarks | Method | Ref. |
|---|---|---|---|---|---|---|---|---|
| 1. | Aβ(12-24) | 13 | 10.60 | 0.05 | 25 | water | PFG-NMR | [20] |
| 2. | Aβ(1-28) | 28 | 14.60 | 0.09 | 25 | water | PFG-NMR | [20] |
| 3. | Aβ(1-40) | 40 | 16.1 | 0.2 | 25 | water | PFG-NMR | [20] |
| 4. | pSic1 | 90 | 19.3 | 1.4 | 25 | 10 mM sodium phosphate pH 7.0, 140 mM NaCl, 1 mM EDTA, 0.2% $NaN_3$, 10% $D_2O$ | PFG-NMR | [27] |
| 5. | p53(1-93) | 93 | 32.8 | 1.3 | 25 | 10 mM sodium phosphate pH 7, 100 mM NaCl | DLS | [31] |
| 6. | $E_m$ protein | 93 | 28 | no data | 23 | 20 mM HEPES pH 7.5, 100 mM KCl, 0.5 mM DTT, 3 mM $MgCl_2$ | SEC | [17] |
| 7. | Lysozyme | 129 | 17 | 1 | 25 | 50 mM Tris pH 8.0, 150 mM NaCl, 0.5 mM EDTA, 1 mM TCEP or DTT globular protein [1)] | FCS | This work |
| 8. | AaFEcR | 131 | 27 | 1 | RT | 10 mM Tris-HCl pH 7.0, 150 mM NaCl | SEC | [34] |
| 9. | Aap PGR | 135 | 38.4 | 0.9 | 25 | 20 mM $K_2HPO_4/KH_2PO_4$ pH 7.4, 150 mM NaCl | DLS | [32] |
| 10. | $N_{TAIL}$ | 139 | 30 | 2 | 20 | 10 mM sodium phosphate pH 7 and in 10 mM Tris pH 8, 75 mM NaCl | DLS | [22] |
| 11. | $\alpha$-Synuclein | 140 | 27.9 | 0.3 | 20 | 20 mM $Na_2HPO_4/NaH_2PO_4$ pH 7.4, 150 mM NaCl, 2% glycerol, 10% $D_2O$, 0.25 mM DSS, 0.02% dioxane, 0.02% $NaN_3$ | PFG-NMR | [35] |
| 12. | hNL3-cyt | 140 | 27.3 | 0.4 | 20 | Phosphate-buffered saline pH 8.0 | AUC | [28] |

| Id. | Protein name | N | $R_h$ exp [Å] | $\Delta R_h$ exp [Å] | T [°C] | Conditions/ Remarks | Method | Ref. |
|---|---|---|---|---|---|---|---|---|
| 13. | SAP 1A | 149 | 33 | 3 | 25 | 50 mM Tris pH 8.0, 150 mM NaCl, 0.5 mM EDTA, 1 mM TCEP or DTT | FCS | This work |
| 14. | $ANAC046_{172-338}$ | 167 | 30.4 | 0.1 | 25 | 20 mM $Na_2HPO_4/NaH_2PO_4$ pH 7.0 100 mM NaCl, 1 mM DTT, 0.02% dioxane, 0.02% $NaN_3$ | PFG-NMR | [35] |
| 15. | HIF-1α (530-698) | 169 | 38.30 | 0.04 | RT | 25 mM NaPi pH 7.2, 150 mM KCl, 10 mM 2-mercaptoethanol | SEC | [24] |
| 16. | eIF4E(28-217) | 190 | 27.0 | 1.9 | 25 | 50 mM Tris pH 8.0, 150 mM NaCl, 0.5 mM EDTA, 1 mM TCEP or DTT | FCS | This work |
| 17. | HIF-1α (403-603) | 201 | 44.30 | 0.1 | RT | 25 mM NaPi pH 7.2, 150 mM KCl, 10 mM 2-mercaptoethanol | SEC | [24] |
| 18. | Securin | 204 | 39.70 | 0.04 | RT | 25 mM NaPi pH 7.2, 150 mM KCl, 10 mM 2-mercaptoethanol | SEC | [23] |
| 19. | SNAP25 | 206 | 39.3 | 2.8 | 20 | 20 mM Tris pH 7.5, 150 mM NaCl, 0.5 mM DTT | DLS | [30] |
| 20. | eIF4E | 217 | 32.4 | 1.7 | 25 | 50 mM Tris pH 8.0, 150 mM NaCl, 0.5 mM EDTA, 1 mM TCEP or DTT | FCS | This work |
| 21. | $H_6$-PNT | 236 | 47 | 4 | 20 | 100 mM NaCl pH 8 | DLS | [21] |
| 22. | 3D7-6H MSP2 | 237 | 34.3 | 0.7 | 25 | PBS pH 7.0 | PFG-NMR | [26] |
| 23. | a-chymo-trypsynogen A | 245 | 23 | 1 | 25 | 50 mM Tris pH 8.0, 150 mM NaCl, 0.5 mM EDTA, 1 mM TCEP or DTT globular protein | FCS | This work |
| 24. | $H_6$-mCherry | 256 | 25.8 | 0.7 | 25 | 50 mM Tris pH 8.0, 150 mM NaCl, 0.5 mM EDTA, 1 mM TCEP or DTT globular protein | FCS | This work |
| 25. | $H_6$-SUMO-SAP 1A | 267 | 37 | 4 | 25 | 50 mM Tris pH 8.0, 150 mM NaCl, 0.5 mM EDTA, 1 mM TCEP or DTT | FCS | This work |
| 26. | maEGFP-$H_6$ | 272 | 22.2 | 0.6 | 25 | 50 mM Tris pH 8.0, 150 mM NaCl, 0.5 mM EDTA, 1 mM TCEP or DTT globular protein | FCS | This work |

| Id. | Protein name | N | $R_h$ exp [Å] | $\Delta R_h$ exp [Å] | T [°C] | Conditions/ Remarks | Method | Ref. |
|---|---|---|---|---|---|---|---|---|
| 27. | $H_6$-mCherry-a-helix | 282 | 30.5 | 0.9 | 25 | 50 mM Tris pH 8.0, 150 mM NaCl, 0.5 mM EDTA, 1 mM TCEP or DTT | FCS | This work |
| 28. | $H_6$-SUMO-CNOT1(800-999) | 324 | 29.6 | 1.2 | 25 | 50 mM Tris pH 8.0, 150 mM NaCl, 0.5 mM EDTA, 1 mM TCEP or DTT | FCS | This work |
| 29. | GW182 SD ΔRRM | 348 | 36 | 3 | 25 | 50 mM Tris pH 8.0, 150 mM NaCl, 0.5 mM EDTA, 1 mM TCEP or DTT | FCS | This work |
| 30. | SUMO-maEGFP-$H_6$ | 370 | 35 | 6 | 25 | 50 mM Tris pH 8.0, 150 mM NaCl, 0.5 mM EDTA, 1 mM TCEP or DTT | FCS | This work |
| 31. | Calreticulin | 404 | 46 | no data | RT | 20 mM Hepes pH 7.5, 150 mM NaCl | SEC | [19] |
| 32. | HeV PNT | 410 | 44 | 2 | RT | 10 mM Tris buffer pH 8, 300 mM NaCl and in 10 mM sodium phosphate pH 7, 150 mM NaCl | SEC | [29] |
| 33. | NiV PNT | 412 | 44 | 2 | RT | 10 mM Tris buffer pH 8, 300 mM NaCl and in 10 mM sodium phosphate pH 7, 150 mM NaCl | SEC | [29] |
| 34. | GST-CNOT1(800-999) | 434 | 39 | 2 | 25 | 50 mM Tris pH 8.0, 150 mM NaCl, 0.5 mM EDTA, 1 mM TCEP or DTT | FCS | This work |
| 35. | $H_6$-SUMO-GW182 SD ΔRRM | 469 | 38 | 3 | 25 | 50 mM Tris pH 8.0, 150 mM NaCl, 0.5 mM EDTA, 1 mM TCEP or DTT | FCS | This work |
| 36. | AGARP | 506 | 54 | 3 | 25 | 50 mM Tris pH 8.0, 150 mM NaCl, 0.5 mM EDTA, 1 mM TCEP or DTT | FCS | This work |
| 37. | $H_6$-SUMO-PARNC-mCherry | 513 | 44.4 | 1.9 | 25 | 50 mM Tris pH 8.0, 150 mM NaCl, 0.5 mM EDTA, 1 mM TCEP or DTT | FCS | This work |
| 38. | HSA | 585 | 33.4 | 1.7 | 25 | 50 mM Tris pH 8.0, 150 mM NaCl, 0.5 mM EDTA, 1 mM TCEP or DTT globular protein | FCS | This work |
| 39. | OMM-64 | 608 | 75.9 | 0.1 | 20 | 10 mM Tris pH 7.5, 100 mM NaCl | AUC | [33] |
| 40. | $H_6$-SUMO-AGARP | 624 | 57 | 3 | 25 | 50 mM Tris pH 8.0, 150 mM NaCl, 0.5 mM EDTA, 1 mM TCEP or DTT | FCS | This work |

| Id. | Protein name | N | $R_h$ exp [Å] | $\Delta R_h$ exp [Å] | T [°C] | Conditions/ Remarks | Method | Ref. |
|---|---|---|---|---|---|---|---|---|
| 41. | GW182SD-mCherry | 688 | 50 | 3 | 25 | 50 mM Tris pH 8.0, 150 mM NaCl, 0.5 mM EDTA, 1 mM TCEP or DTT | FCS | This work |
| 42. | $H_6$-SUMO-GW182 SD-mCherry | 809 | 66 | 6 | 25 | 50 mM Tris pH 8.0, 150 mM NaCl, 0.5 mM EDTA, 1 mM TCEP or DTT | FCS | This work |
| 43. | Fesselin | 996 | 53 | no data | RT? | 20 mM MOPS pH 7.0, 200 mM NaCl, 2 mM EGTA | SEC | [25] |
| 44. | Apoferritin (24-mer) | 4200 | 58 | 3 | 25 | 50 mM Tris pH 8.0, 150 mM NaCl, 0.5 mM EDTA, 1 mM TCEP or DTT globular protein [2)] | FCS | This work |

[1)] the $R_h$ value from FCS is slightly underestimated due to the residual presence of the freely diffusing dye impossible to be completely separated from the protein by SEC and the short diffusion time of lysozyme.

[2)] shown in Figures 2 and 3A (main text) for comparison with other proteins; not included in the analysis of the theoretical model

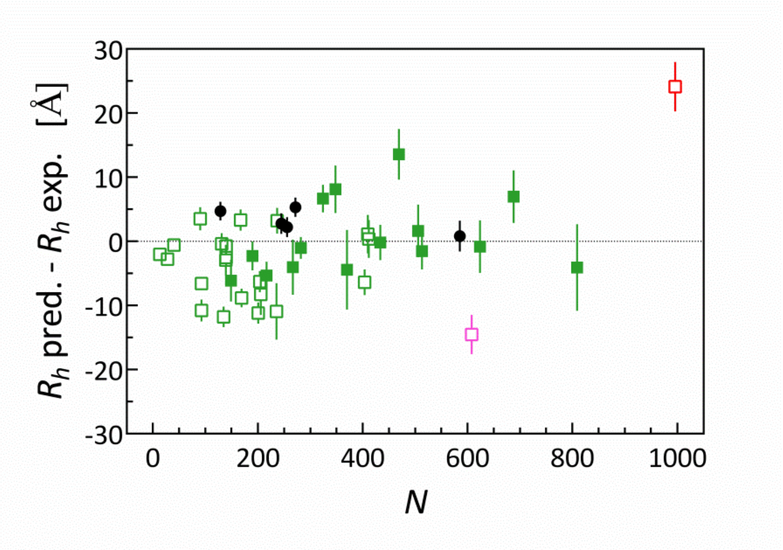


**Figure S7.** Difference between $R_h$ values predicted by MDA+GLM (Table S2) and experimental results (Table S1) for the benchmark set. IDPs (full green squares) and folded proteins (full black circles) from this work; IDPs from literature (blank squares); two largest outliers are marked in red (fesselin, Id. 43, N = 996, SEC) and magenta (OMM-64, Id. 39, N = 608, AUC). Error bars reflect both theoretical (Table S2, column F) and experimental uncertainties (Table S1) calculated according to small errors propagation rules.

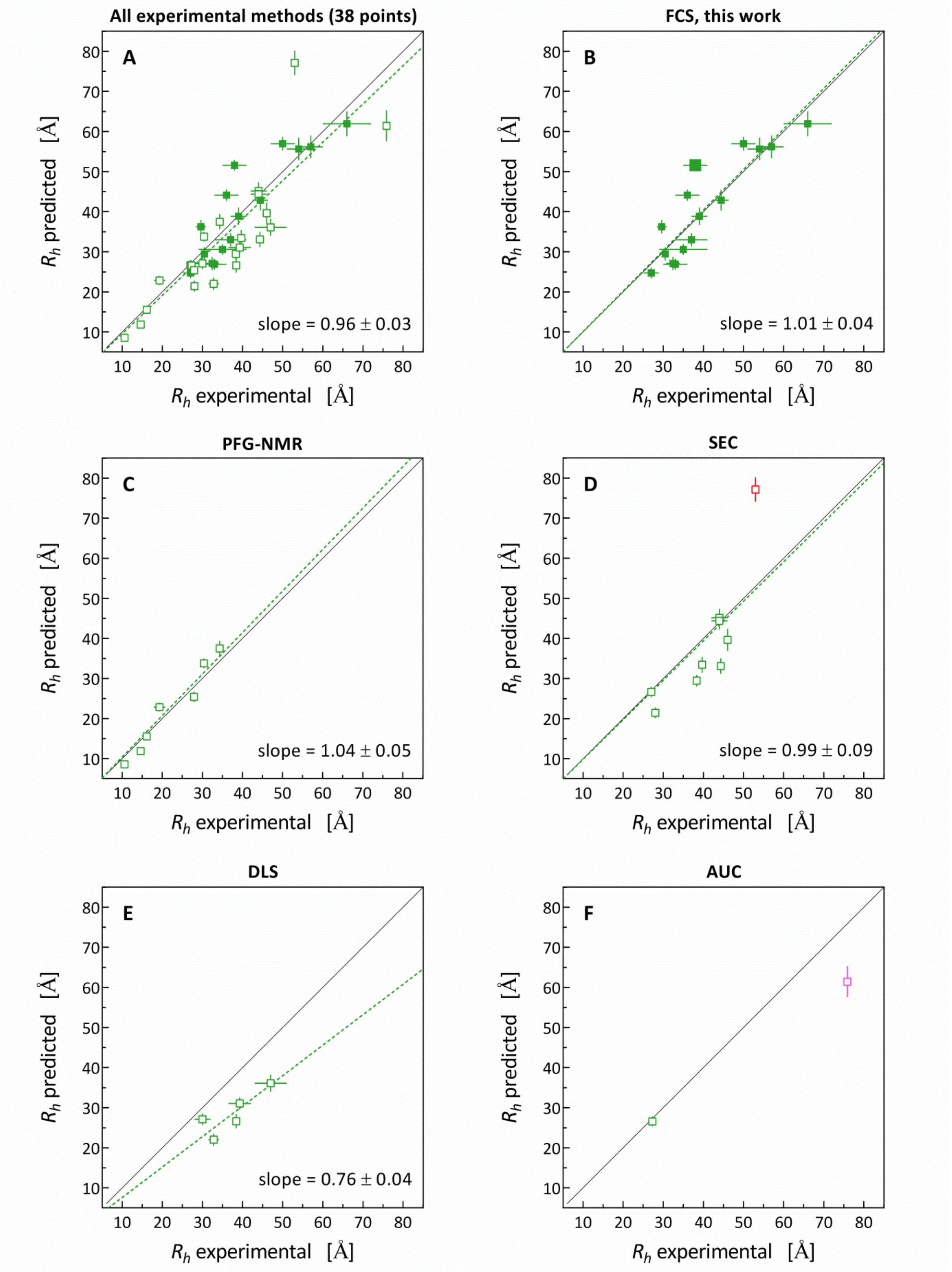


**Figure S8.** Correlation analysis of $R_h$ values predicted for IDPs by MDA+GLM (Table S2, columns D, F) *vs.* experimental results (Table S1, the benchmark set excluding globular proteins); 1:1 relationship (thin black line); linear fit to the data points without free *y*-intercept (green broken line, except F); (A) all $R_h$ values (full green squares, this work; blank green squares, literature); (B-F) subsets of results obtained using different experimental approaches, *i.e.* PFG-NMR, FCS (this work), SEC, DLS, and AUC, respectively.

The Snedecor's *F*-test for the linear functions with and without the *y*-intercept as a free parameter fitted to the data points from the IDPs benchmark set showed that the *y*-intercept is insignificantly different from zero, -0.26 ± 3.6. The fit (**Figure S8 A**) yielded the slope of 0.96 ± 0.03 (with 90% confidence interval, CI, of 0.905 to 1.006). This means that MDA+GLM provides good 1:1 correlation with the experimental results for IDPs even at the level of 90% CI.

The $R^2$ of the linear correlation between the predicted and experimental results for all IDPs from Table S1 is 0.7534 (**Figure S8 A)**, which means that our model explains ~75% of the $R_h$ variability within the IDP benchmark set. The remaining part of the variability as well as the slightly underestimated slope value can have several sources. Among the main reasons for the discrepancies are the intrinsic properties of individual experimental methods, which suffer from typical errors or limitations and are usually not taken into account when reporting the final experimental results.

The root mean square of the relative uncertainty for all experimental data (Table S1), when given, is 5.8%. Even for a perfect model that accurately predicts the diffusion coefficient, assuming the measurement uncertainty is only random (not systematic), achieving $R^2 = 1$ is impossible due to the inherent random noise in the data. The median $R^2$ values under these conditions, determined theoretically, are gathered in Table S5.

**Table S5.**

| Relative error % | Median $R^2$ of a perfect model |
|---|---|
| 5 | 0.98 |
| 5.8 | 0.97 |
| 10 | 0.92 |
| 15 | 0.85 |
| 20 | 0.76 |

However, the value of 5.8% seems underestimated. This is because it relies on undervalued figures provided in literature, where only some parts of the uncertainty are included in the error estimates, and in some cases, no error analysis is provided. Assuming a more realistic overall measurement error of 10%, which may still be considered small for certain measurements, the best possible model should give a typical $R^2$ of ~0.9.

Considering that our GLM-MDA approach involves approximated hydrodynamics, the predictions result in ~5% error of the theoretical $R_h$ values. Therefore, one should expect results only up to an $R^2$ of 0.85, even with exceptionally precise modeling of conformers, hydration layers, and other complex factors.

**Intrinsically disordered benchmark proteins gathered in Table S1.**
Sequence numbering according to Table S1.

**Data of protein constructs studied in this work**
Fusion proteins may contain linkers between the domains identified in the protein names
Abbreviations used repeatedly in protein names:

| | | |
|---|---|---|
| $H_6$ | - | Hexahistidine tag |
| SUMO | - | Small ubiquitin-related modifier protein tag |
| GST | - | Glutathione S-transferase tag |
| mCherry | - | Monomeric red fluorescent protein[36] |
| mαEGFP | - | Enhanced green fluorescent protein[37] with mutations providing the monomeric form[38] and improved by α mutations[39] |

**13. SAP 1A** (Secreted acidic protein 1A, *Acropora millepora*, UniProtKB B3EWZ0)
GLPLPLKNENAIVDGDGTSVVTTKEDASTIFERDPNPANQVSAMVTGVILDENGDPGESDESVENVDNDGEGGDK
DDDKNGEDNDLDNKEHEEEKGDDDRGDDEEEDDAEGDNDSNDNEGDDDDDDDSGDDDDVDESGADEDDDDDSGD

**16. eIF4E(28-217)** (Eukaryotic translation initiation 4e factor, *Mus musculus*, UniProtKB P63073)
VANPEHYIKHPLQNRWALWFFKNDKSKTWQANLRLISKFDTVEDFWALYNHIQLSSNLMPGCDYSLFKDGIEPMW
EDEKNKRGGRWLITLNKQQRRSDLDRFWLETLLCLIGESFDDYSDDVCGAVVNVRAKGDKIAIWTTECENRDAVT
HIGRVYKERLGLPPKIVIGYQSHADTATKSGSTTKNRFVV

**20. eIF4E** (Eukaryotic translation initiation 4e factor, *Homo sapiens*, UniProtKB P06730)
MATVEPETTPTPNPPTTEEEKTESNQEVANPEHYIKHPLQNRWALWFFKNDKSKTWQANLRLISKFDTVEDFWAL
YNHIQLSSNLMPGCDYSLFKDGIEPMWEDEKNKRGGRWLITLNKQQRRSDLDRFWLETLLCLIGESFDDYSDDVC
GAVVNVRAKGDKIAIWTTECENREAVTHIGRVYKERLGLPPKIVIGYQSHADTATKSGSTTKNRFVV

**25. $H_6$-SUMO-SAP 1A**
MGSSHHHHHHSSGLVPRGSHMSDSEVNQEAKPEVKPEVKPETHINLKVSDGSSEIFFKIKKTTPLRRLMEAFAKR
QGKEMDSLRFLYDGIRIQADQTPEDLDMEDNDIIEAHREQIGGGLPLPLKNENAIVDGDGTSVVTTKEDASTIFE
RDPNPANQVSAMVTGVILDENGDPGESDESVENVDNDGEGGDKDDDKNGEDNDLDNKEHEEEKGDDDRGDDEEED
DAEGDNDSNDNEGDDDDDDDSGDDDDVDESGADEDDDDDSGD

**27. $H_6$-mCherry-α-helix**
MGSSHHHHHHSSGLVPRGSHMVSKGEEDNMAIIKEFMRFKVHMEGSVNGHEFEIEGEGEGRPYEGTQTAKLKVTK
GGPLPFAWDILSPQF***MYG***SKAYVKHPADIPDYLKLSFPEGFKWERVMNFEDGGVVTVTQDSSLQDGEFIYKVKLR
GTNFPSDGPVMQKKTMGWEASSERMYPEDGALKGEIKQRLKLKDGGHYDAEVKTTYKAKKPVQLPGAYNVNIKLD
ITSHNEDYTIVEQYERAEGRHSTGGMDELYKGTGVDQDPAANKARKEAELAAATAQQ

**28. $H_6$-SUMO-CNOT1(800-999)** (CCR4-NOT transcription complex subunit 1, isoform 2, *Homo sapiens*, UniProtKB A5YKK6)
MGSSHHHHHHSSGLVPRGSHMASMSDSEVNQEAKPEVKPEVKPETHINLKVSDGSSEIFFKIKKTTPLRRLMEAF
AKRQGKEMDSLRFLYDGIRIQADQTPEDLDMEDNDIIEAHREQIGGSEFNNDPFVQRKLGTSGLNQPTFQQTDLS
QVWPEANQHFSKEIDDEANSYFQRIYNHPPHPTMSVDEVLEMLQRFKDSTIKREREVFNCMLRNLFEEYRFFPQY
PDKELHITACLFGGIIEKGLVTYMALGLALRYVLEALRKPFGSKMYYFGIAALDRFKNRLKDYPQYCQHLASISH
FMQFPHHLQEYIEYGQQSRDPPVK

**29. GW182 SD ΔRRM** (Silencing domain of trinucleotide repeat-containing gene 6C protein, without the globular RRM domain, *Homo sapiens*, UniProtKB Q9HCJ0)
SEFNTFAPYPLAGLNPNMNVNSMDMTGGLSVKDPSQSQSRLPQWTHPNSMDNLPSAASPLEQNPSKHGAIPGGLS
IGPPGKSSIDDSYGRYDLIQNSESPASPPVAVPHSWSRAKSDSDKISNGSSINWPPEFHPGVPWKGLQNIDPEND
PDVTPGSVPTGPTINTTIQDVNRYLLKSGGKLSDIKSTWSSGPTSHTQASLSHELWKVPRNSTAPTRPPPGLTNP
KPSSTWGASPLGWTSSYSSGSAWSTDTSGQALPPTSSWQSSSASSQPRLSAAGSSHGLVRSDAGHWNAPCLGGKG
SSELLWGGVPQYSSSLWGPPSADDSRVIGSPTPLTTLLPGDLLSGESL

**30. SUMO-mαEGFP-$H_6$**
MSDSEVNQEAKPEVKPEVKPETHINLKVSDGSSEIFFKIKKTTPLRRLMEAFAKRQGKEMDSLRFLYDGIRIQAD
QTPEDLDMEDNDIIEAHREQIGGGTVSKGEELFTGVVPILVELDGDVNGHKFSVSGEGEGDATYGKLTLKFICTT
GKLPVPWPTLVTTL***TYG***VQCFSRYPDHMKRHDFFKSAMPEGYVQERTISFKDDGNYKTRAEVKFEGDTLVNRIEL
KGIDFKEDGNILGHKLEYNYNSHNVYITADKQKNGIKANFKIRHNIEDGSVQLADHYQQNTPIGDGPVLLPDNHY
LSTQSKLSKDPNEKRDHMVLLEFVTAAGITLGMDELYKDIGSAGSAAGSGEFEFLEVLFQGPLEHHHHHH

**34. GST-CNOT1(800-999)**
MSPILGYWKIKGLVQPTRLLLEYLEEKYEEHLYERDEGDKWRNKKFELGLEFPNLPYYIDGDVKLTQSMAIIRYI
ADKHNMLGGCPKERAEISMLEGAVLDIRYGVSRIAYSKDFETLKVDFLSKLPEMLKMFEDRLCHKTYLNGDHVTH
PDFMLYDALDVVLYMDPMCLDAFPKLVCFKKRIEAIPQIDKYLKSSKYIAWPLQGWQATFGGGDHPPKSDLEVLF
QGPLGSPEFNNDPFVQRKLGTSGLNQPTFQQTDLSQVWPEANQHFSKEIDDEANSYFQRIYNHPPHPTMSVDEVL
EMLQRFKDSTIKREREVFNCMLRNLFEEYRFFPQYPDKELHITACLFGGIIEKGLVTYMALGLALRYVLEALRKP
FGSKMYYFGIAALDRFKNRLKDYPQYCQHLASISHFMQFPHHLQEYIEYGQQSRDPPVK

**35. $H_6$-SUMO-GW182 SD ΔRRM**
MGSSHHHHHHSSGLVPRGSHMASMSDSEVNQEAKPEVKPEVKPETHINLKVSDGSSEIFFKIKKTTPLRRLMEAF
AKRQGKEMDSLRFLYDGIRIQADQTPEDLDMEDNDIIEAHREQIGGSEFNTFAPYPLAGLNPNMNVNSMDMTGGL
SVKDPSQSQSRLPQWTHPNSMDNLPSAASPLEQNPSKHGAIPGGLSIGPPGKSSIDDSYGRYDLIQNSESPASPP
VAVPHSWSRAKSDSDKISNGSSINWPPEFHPGVPWKGLQNIDPENDPDVTPGSVPTGPTINTTIQDVNRYLLKSG
GKLSDIKSTWSSGPTSHTQASLSHELWKVPRNSTAPTRPPPGLTNPKPSSTWGASPLGWTSSYSSGSAWSTDTSG
QALPPTSSWQSSSASSQPRLSAAGSSHGLVRSDAGHWNAPCLGGKGSSELLWGGVPQYSSSLWGPPSADDSRVIG
SPTPLTTLLPGDLLSGESL

**36. AGARP** (Aspartic and glutamic acid-rich protein, *Acropora millepora*, UniProtKB B7W112)
SPLRNRFNEDHDEFSKDDMARESFDTEEMYNAFLNRRDSSESQLEDHLLSHAKPLYDDFFPKDTSPDDDEDSYWL
ESRNDDGYDLAKRKRGYDDEEAYDDFDEVDDRADDEGARDVDESDFEEDDKLPAEEESKNDMDEETFEDEPEEDK
EEAREEFAEDERADEREDDDADFDFNDEEDEDEVDNKAESDIFTPEDFAGVSDEAMDNFRDDNEEEYADESDDEA
EEDSEETADDFEDDPEDESDETFRDEVEDESEENYQDDTEEGSEIKQNDETEEQPEKKFDADKEHEDAPEPLKEK
LSDESKARAEDESDKSEDAAKEIKEPEDAVEDFEDGAKVSEDEAELLDDEAELSDDEAELSKDEAEQSSDEAEKS
EDKAEKSEDEAELSEDEAKQSEDEAEKAEDAAGKESNDEGKKREDEAVKSKGIARDESEFAKAKKSNLALKRDEN
RPLAKGLRESAAHLRDFPSEKKSKDAAQGNIENELDYFKRNAFADSKDAEPYEFDK

**37. $H_6$-SUMO-PARN C-mCherry** (Poly(A)-specific ribonuclease C-terminal tail, *Homo sapiens*, UniProtKB O95453)
MGSSHHHHHHSSGLVPRGSHMSDSEVNQEAKPEVKPEVKPETHINLKVSDGSSEIFFKIKKTTPLRRLMEAFAKR
QGKEMDSLRFLYDGIRIQADQTPEDLDMEDNDIIEAHREQIGGYAESYRIQTYAEYMGRKQEEKQIKRKWTEDSW
KEADSKRLNPQCIPYTLQNHYYRNNSFTAPSTVGKRNLSPSQEEAGLEDGVSGEISDTELEQTDSCAEPLSEGRK
KAKKLKRMKKELSPAGSISKNSPATLFEVPDTWLEVLFQGPGSAGSAAGSGEFVSKGEEDNMAIIKEFMRFKVHM
EGSVNGHEFEIEGEGEGRPYEGTQTAKLKVTKGGPLPFAWDILSPQFMYGSKAYVKHPADIPDYLKLSFPEGFKW
ERVMNFEDGGVVTVTQDSSLQDGEFIYKVKLRGTNFPSDGPVMQKKTMGWEASSERMYPEDGALKGEIKQRLKLK
DGGHYDAEVKTTYKAKKPVQLPGAYNVNIKLDITSHNEDYTIVEQYERAEGRHSTGGMDELYK

**40. $H_6$-SUMO-AGARP**
MGSSHHHHHHSSGLVPRGSHMSDSEVNQEAKPEVKPEVKPETHINLKVSDGSSEIFFKIKKTTPLRRLMEAFAKR
QGKEMDSLRFLYDGIRIQADQTPEDLDMEDNDIIEAHREQIGGSPLRNRFNEDHDEFSKDDMARESFDTEEMYNA
FLNRRDSSESQLEDHLLSHAKPLYDDFFPKDTSPDDDEDSYWLESRNDDGYDLAKRKRGYDDEEAYDDFDEVDDR
ADDEGARDVDESDFEEDDKLPAEEESKNDMDEETFEDEPEEDKEEAREEFAEDERADEREDDDADFDFNDEEDED
EVDNKAESDIFTPEDFAGVSDEAMDNFRDDNEEEYADESDDEAEEDSEETADDFEDDPEDESDETFRDEVEDESE
ENYQDDTEEGSEIKQNDETEEQPEKKFDADKEHEDAPEPLKEKLSDESKARAEDESDKSEDAAKEIKEPEDAVED
FEDGAKVSEDEAELLDDEAELSDDEAELSKDEAEQSSDEAEKSEDKAEKSEDEAELSEDEAKQSEDEAEKAEDAA
GKESNDEGKKREDEAVKSKGIARDESEFAKAKKSNLALKRDENRPLAKGLRESAAHLRDFPSEKKSKDAAQGNIE
NELDYFKRNAFADSKDAEPYEFDK

**41. GW182 SD-mCherry**
SEFNTFAPYPLAGLNPNMNVNSMDMTGGLSVKDPSQSQSRLPQWTHPNSMDNLPSAASPLEQNPSKHGAIPGGLS
IGPPGKSSIDDSYGRYDLIQNSESPASPPVAVPHSWSRAKSDSDKISNGSSINWPPEFHPGVPWKGLQNIDPEND
PDVTPGSVPTGPTINTTIQDVNRYLLKSGGKLSDIKSTWSSGPTSHTQASLSHELWKVPRNSTAPTRPPPGLTNP
KPSSTWGASPLGWTSSYSSGSAWSTDTSGRTSSWLVLRNLTPQIDGSTLRTLCLQHGPLITFHLNLTQGNAVVRY

SSKEEAAKAQKSLHMCVLGNTTILAEFAGEEEVNRFLAQGQALPPTSSWQSSSASSQPRLSAAGSSHGLVRSDAG
HWNAPCLGGKGSSELLWGGVPQYSSSLWGPPSADDSRVIGSPTPLTTLLPGDLLSGESLPGGSAGSAAGSGEFAA
AVSKGEEDNMAIIKEFMRFKVHMEGSVNGHEFEIEGEGEGRPYEGTQTAKLKVTKGGPLPFAWDILSPQF***MYG***SK
AYVKHPADIPDYLKLSFPEGFKWERVMNFEDGGVVTVTQDSSLQDGEFIYKVKLRGTNFPSDGPVMQKKTMGWEA
SSERMYPEDGALKGEIKQRLKLKDGGHYDAEVKTTYKAKKPVQLPGAYNVNIKLDITSHNEDYTIVEQYERAEGR
HSTGGMDELYKKL

**42. $H_6$-SUMO-GW182 SD-mCherry**
MGSSHHHHHHSSGLVPRGSHMASMSDSEVNQEAKPEVKPEVKPETHINLKVSDGSSEIFFKIKKTTPLRRLMEAF
AKRQGKEMDSLRFLYDGIRIQADQTPEDLDMEDNDIIEAHREQIGGSEFNTFAPYPLAGLNPNMNVNSMDMTGGL
SVKDPSQSQSRLPQWTHPNSMDNLPSAASPLEQNPSKHGAIPGGLSIGPPGKSSIDDSYGRYDLIQNSESPASPP
VAVPHSWSRAKSDSDKISNGSSINWPPEFHPGVPWKGLQNIDPENDPDVTPGSVPTGPTINTTIQDVNRYLLKSG
GKLSDIKSTWSSGPTSHTQASLSHELWKVPRNSTAPTRPPPGLTNPKPSSTWGASPLGWTSSYSSGSAWSTDTSG
RTSSWLVLRNLTPQIDGSTLRTLCLQHGPLITFHLNLTQGNAVVRYSSKEEAAKAQKSLHMCVLGNTTILAEFAG
EEEVNRFLAQGQALPPTSSWQSSSASSQPRLSAAGSSHGLVRSDAGHWNAPCLGGKGSSELLWGGVPQYSSSLWG
PPSADDSRVIGSPTPLTTLLPGDLLSGESLPGGSAGSAAGSGEFAAAVSKGEEDNMAIIKEFMRFKVHMEGSVNG
HEFEIEGEGEGRPYEGTQTAKLKVTKGGPLPFAWDILSPQF***MYG***SKAYVKHPADIPDYLKLSFPEGFKWERVMNF
EDGGVVTVTQDSSLQDGEFIYKVKLRGTNFPSDGPVMQKKTMGWEASSERMYPEDGALKGEIKQRLKLKDGGHYD
AEVKTTYKAKKPVQLPGAYNVNIKLDITSHNEDYTIVEQYERAEGRHSTGGMDELYKKL

#### Literature data

**1. Aβ(12–24)**
VHHQKLVFFAEDV

**2. Aβ(1–28)**
DAEFRHDSGYEVHHQKLVFFAEDVGSNK

**3. Aβ(1–40)**
DAEFRHDSGYEVHHQKLVFFAEDVGSNKGAIIGLMVGGVV

**4. pSic1**
MTPSTPPRSRGTRYLAQPSGNTSSSALMQGQKTPQKPSQNLVPVTPSTTKSFKNAPLLAPPNSNMGMTSPFNGLT
SPQRSPFPKSSVKRT

**5. p53(1-93)**
MEEPQSDPSVEPPLSQETFSDLWKLLPENNVLSPLPSQAMDDLMLSPDDIEQWFTEDPGPDEAPRMPEAAPPVAP
APAAPTPAAPAPAPSWPL

**6. $E_m$ protein**
MASGQQERSQLDRKAREGETVVPGGTGGKSLEAQENLAEGRSRGGQTRREQMGEEGYSQMGRKGGLSTNDESGGD
RAAREGIDIDESKFKTKS

**8. AaFEcR**
GPSAGLVPRGSGGIEGRHMLEEIWDVQDIPPSMQAQMHSHGTQSSSSSSSSSSSSSNGSSNGNSSSNSNSNSSQHGP
HPHPHGQQLTPNQQQHQQQHSQLQQVHANGSGSGGGSNNNSSSGGVVPGLGMLDQV

**9. Aap PGR**
AEPGKPAEPGKPAEPGKPAEPGTPAEPGKPAEPGTPAEPGKPAEPGKPAEPGKPAEPGKPAEPGTPAEPGTPAEP
GKPAEPGTPAEPGKPAEPGTPAEPGKPAESGKPVEPGTPAQSGAPEQPNRSMHSTDNKNQ

**10. $N_{TAIL}$**
MRGSHHHHHHXXXHTTEDKISRAVGPRQAQVSFLHGDQSENELPRLGGKEDRRVKQSRGEARESYRETGPSRASD
ARAAHLPTGTPLDIDTASESSQDPQDSRRSADALLRLQAMAGISEEQGSDTDTPIVYNDRNLLD

**11. α-Synuclein**
MDVFMKGLSKAKEGVVAAAEKTKQGVAEAAGKTKEGVLYVGSKTKEGVVHGVATVAEKTKEQVTNVGGAVVTGVT
AVAQKTVEGAGSIAAATGFVKKDQLGKNEEGAPQEGILEDMPVDPDNEAYEMPSEEGYQDYEPEA

**12. hNL3-cyt**
MGSSHHHHHHSSGLVPRGSHMAYRKDKRRQEPLRQPSPQRGAGAPELGAAPEEELAALQLGPTHHECEAGPPHDT
LRLTALPDYTLTLRRSPDDIPLMTPNTITMIPNSLVGLQTLHPYNTFAAGFNSTGLPHSHSTTRV

**14. ANAC046$_{172\text{-}338}$**
NAPSTTITTTKQLSRIDSLDNIDHLLDFSSLPPLIDPGFLGQPGPSFSGARQQHDLKPVLHHPTTAPVDNTYLPT
QALNFPYHSVHNSGSDFGYGAGSGNNNKGMIKLEHSLVSVSQETGLSSDVNTTATPEISSYPMMMNPAMMDGSKS
ACDGLDDLIFWEDLYTS

**15. HIF-1α (530-698)**
XEFKLELVEKLFAEDTEAKNPFSTQDTDLDLEMLAPYIPMDDDFQLRSFDQLSPLESSSASPESASPQSTVTVFQ
QTQIQEPTANATTTTATTDELKTVTKDRMEDIKILIASPSPTHIHKETTSATSSPYRDTQSRTASPNRAGKGVIE
QTEKSHPRSPNVLSVALSQR

**17. HIF-1α (403-603)**
AAGDTIISLDFGSNDTETDDQQLEEVPLYNDVMLPSPNEKLQNINLAMSPLPTAETPKPLRSSADPALNQEVALK
LEPNPESLELSFTMPQIQDQTPSPSDGSTRQSSPEPNSPSEYCFYVDSDMVNEFKLELVEKLFAEDTEAKNPFST
QDTDLDLEMLAPYIPMDDDFQLRSFDQLSPLESSSASPESASPQSTVTVFQ

**18. Securin**
XXMATLIYVDKENGEPGTRVVAKDGLKLGSGPSIKALDGRSQVSTPRFGKTFDAPPALPKATRKALGTVNRATEK
SVKTKGPLKQKQPSFSAKKMTEKTVKAKSSVPASDDAYPEIEKFFPFNPLDFESFDLPEEHQIAHLPLSGVPLMI
LDEERELEKLFQLGPPSPVKMPSPPWESNLLQSPSSILSTLDVELPPVCCDIDI

**19. SNAP25**
MAEDADMRNELEEMQRRADQLADESLESTRRMLQLVEESKDAGIRTLVMLDEQGEQLERIEEGMDQINKDMKEAE
KNLTDLGKFCGLCVCPCNKLKSSDAYKKAWGNNQDGVVASQPARVVDEREQMAISGGFIRRVTNDARENEMDENL
EQVSGIIGNLRHMALDMGNEIDTQNRQIDRIMEKADSNKTRIDEANQRATKMLGSG

**21. $H_6$-PNT**
HHHHHHMAEEQARHVKNGLECIRALKAEPIGSLAIEEAMAAWSEISDNPGQERATCREEKAGSSGLSKPCLSAIG
STEGGAPRIRGQGPGESDDDAETLGIPPRNLQASSTGLQCYYVYDHSGEAVKGIQDADSIMVQSGLDGDSTLSGG
DNESENSDVDIGEPDTEGYAITDRGSAPISMGFRASDVETAEGGEIHELLRLQSRGNNFPKLGKTLNVPPPPDPG
RASTSGTPIKK

**22. 3D7-6HMSP2**
MIKNESKYSNTFINNAYNMSIRRSMAESKPSTGAGGSAGGSAGGSAGGSAGGSAGGSAGSGDGNGADAEGSSSTP
ATTTTTKTTTTTTTTNDAEASTSTSSENPNHKNAETNPKGKGEVQEPNQANKETQNNSNVQQDSQTKSNVPPTQD
ADTKSPTAQPEQAENSAPTAEQTESPELQSAPENKGTGQHGHMHGSRNNHPQNTSDSQKECTDGNKENCGAATSL
LNNSSNHHHHHH

**31. Calreticulin**
GIPGEPAVYFKEQFLDGDGWTSRWIESKHKSDFGKFVLSSGKFYGDEEKDKGLQTSQDARFYALSASFEPFSNKG
QTLVVQFTVKHEQNIDCGGGYVKLFPNSLDQTDMHGDSEYNIMFGPDICGPGTKKVHVIFNYKGKNVLINKDIRC
KDDEFTHLYTLIVRPDNTYEVKIDNSQVESGSLEDDWDFLPPKKIKDPDASKPEDWDERAKIDDPTDSKPEDWDK
PEHIPDPDAKKPEDWDEEMDGEWEPPVIQNPEYKGEWKPRQIDNPDYKGTWIHPEIDNPEYSPDPSIYAYDNFGV
LGLDLWQVKSGTIFDNFLITNDEAYAEEFGNETWGVTKAAEKQMKDKQDEEQRLKEEEEDKKRKEEEEAEDKEDD
EDKDEDEEDEEDKEEDEEEDVPGQAKDEL

**32. HeVPNT**
MDKLDLVNDGLDIIDFIQKNQKEIQKTYGRSSIQQPSTKDRTRAWEDFLQSTSGEHEQAEGGMPKNDGGTEGRNV
EDLSSVTSSDGTIGQRVSNTRAWAEDPDDIQLDPMVTDVVYHDHGGECTGHGPSSSPERGWSYHMSGTHDGNVRA
VPDTKVLPNAPKTTVPEEVREIDLIGLEDKFASAGLNPAAVPFVPKNQSTPTEEPPVIPEYYYGSGRRGDLSKSP
PRGNVNLDSIKIYTSDDEDENQLEYEDEFAKSSSEVVIDTTPEDNDSINQEEVVGDPSDQGLEHPFPLGKFPEKE
ETPDVRRKDSLMQDSCKRGGVPKRLPMLSEEFECSGSDDPIIQELEREGSHPGGSLRLREPPQSSGNSRNQPDRQ
LKTGDAASPGGVQRPGTPMPKSRIMPIKKHHHHHH

**33. NiVPNT**

MDKLELVNDGLNIIDFIQKNQKEIQKTYGRSSIQQPSIKDQTKAWEDFLQCTSGESEQVEGGMSKDDGDVERRNL
EDLSSTSPTDGTIGKRVSNTRDWAEGSDDIQLDPVVTDVVYHDHGGECTGYGFTSSPERGWSDYTSGANNGNVCL
VSDAKMLSYAPEIAVSKEDRETDLVHLENKLSTTGLNPTAVPFTLRNLSDPAKDSPVIAEHYYGLGVKEQNVGPQ
TSRNVNLDSIKLYTSDDEEADQLEFEDEFAGSSSEVIVGISPEDEEPSSVGGKPNESIGRTIEGQSIRDNLQAKD
NKSTDVPGAGPKDSAVKEEPPQKRLPMLAEEFECSGSEDPIIRELLKENSLINCQQGKDAQPPYHWSIERSISPD
KTEIVNGAVQTADRQRPGTPMPKSRGIPIKKHHHHHH

**39. OMM-64**

APVNDGTEADNDERAASLLVHLKGDKDGGGLTGSPDGVSAGTTDGTDSSKELAGGAVDSSPDTTDTPDASSSDIF
PDTNNRDTSVETTGNPDDSDAPDAAESAGSQDTTDAADASEAVAETVDTYDIPDTDGADDREKVSTEVSTEDLDS
AGVDKSPESDSTESPGSDSAESPGSDSAESPGSDSTESPGSDSTESPRSDSTDEVLTDVQADSADVTSDDMDEAT
ETDKDDDKSDDKSDADAATDKDDSDEDKDTELDGKAHAEDTQTEEAADSDSKQGAADSDSDTDDDRPEKDVKNDS
DDSKDTTEDDKPDKDDKKNRDSADNSNDDSDEMIQVPREELEQQEINLKEGGVIGSQEETVASDMEEGSDVGDQK
PGPEDSIEEGSPVGRQDFKHPQDSEEEELEKEAKKEKELEEAEEERTLKTIESDSQEDSVDESEAEPDSNSKKDI
GTSDAPEPQEDDSEEDTDDSMMKEPKDSDDAESDKDDKDKNDMDKEDMDKDDMDKDDMDKDDMDKDDVDKDASDS
VDDQSESDAEPGADSHTVVDEIDGEETMTPDSEEIMKSGEMDSVVEATEVPADILDQPDQQDDMTQGASQAADAA
ATALAAQS

**43. Fesselin**

MIQSAAPSIPRVEVILDCSDREKEAPKSLAERGCVDSQVEGGQSEAPPSLPSFAISSEGTEQGEDNQHSEKDHRP
LKHRARHARLRRSESLSEKQVKEAKSKCKSIALLLTAAPNPNSKGVLMFKKRRQRARKYTLVSYGTGELERDEDE
GEEGEVEEGDKENTFEVSLLATSESEIDEDFFSDIDNDKKIVTFDWDSGLLEVEKKTKSGDEMQTLPETTGKGAL
MFARRRQRMDQITAEQEEMKARTAHAEEQREVTVSENFQKVSSSAYQTKEEEMLRQQPCISKSYADVSQNDGKIV
QQNGFGVAPDTSLSFQSSEAQKAASLNRTAKPFPFGVQNRAAAPFSPTRNVTSPLSDLPAPPPYCSISPPPEALY
RPLSAPAASKAAPILWSHTEPTERIASRDERIAVPAKRTGILQEAKRRSTSKPMFSFKEAPKVSPNPALLSLVHN
AEGKKGSGAGFESGPEEDYLSLGAEACNFMQSQASKQKAPPPTAPKPSLKVSPAAGTPVSPVWSPAVASNKAPSF
PAPASPQAAYPAPLKSPQYPHSPSANPPNTLNLSGPFKGPQATLASPNHPAKTPTTPSAGETKPFEMPPEMRGKG
AQLFARRHSRMEKYVVDSETVQANMARASSPTPSLPASWKYSSNVRAPPPVAYNPIHSPSYPPAATKPFPKSTAA
TKNTKRKPKKGLNALDIMKHQPYQLDASLFTFQPPSNKESLGIKQIPKLPTSKQATSLRLPGSASPTNVRASSVY
SVPAYSSQPSFQSNASTPVNESYTPTGYSAFSKPESTTSSLFTAPRPKFSAKKAGVIAQERSSGRSLSLPGKPSF
ISRATSPTSPLIFQPAPDYFSKPDTAADKPGKRLTPWEAAAKSPLGLVDEAFRPQNMQESIAANVVSAAHRKTLP
EPPDEWKQKVSYEPPGPSASLALLGGKQPGVTSARKSSLSVSNATTQAGSQQQYAYCSQRSQTDPDIMSMDSRSD
YGLSTADSNYNPQPRGWRRPT

(13) Jumper, J.; Evans, R.; Pritzel, A.; Green, T.; Figurnov, M.; Ronneberger, O.; Tunyasuvunakool, K.; Bates, R.; Žídek, A.; Potapenko, A.; Bridgland, A.; Meyer, C.; Kohl, S. A. A.; Ballard, A. J.; Cowie, A.; Romera-Paredes, B.; Nikolov, S.; Jain, R.;

Adler, J.; Back, T.; Petersen, S.; Reiman, D.; Clancy, E.; Zielinski, M.; Steinegger, M.; Pacholska, M.; Berghammer, T.; Bodenstein, S.; Silver, D.; Vinyals, O.; Senior, A. W.; Kavukcuoglu, K.; Kohli, P.; Hassabis, D. Highly Accurate Protein Structure Prediction with AlphaFold. *Nature* **2021**, *596* (7873), 583–589. https://doi.org/10.1038/s41586-021-03819-2.
(14) Mirdita, M.; Schütze, K.; Moriwaki, Y.; Heo, L.; Ovchinnikov, S.; Steinegger, M. ColabFold: Making Protein Folding Accessible to All. *Nat Methods* **2022**, *19* (6), 679–682. https://doi.org/10.1038/s41592-022-01488-1.
(15) Jones, D. T.; Cozzetto, D. DISOPRED3: Precise Disordered Region Predictions with Annotated Protein-Binding Activity. *Bioinformatics* **2015**, *31* (6), 857–863. https://doi.org/10.1093/bioinformatics/btu744.
(16) Choi, Y.; Agarwal, S.; Deane, C. M. How Long Is a Piece of Loop? *PeerJ* **2013**, *1*, e1. https://doi.org/10.7717/peerj.1.
(17) McCubbin, W. D.; Kay, C. M.; Lane, B. G. Hydrodynamic and Optical Properties of the Wheat Germ Em Protein. *Can. J. Biochem. Cell Biol.* **1985**, *63* (8), 803–811. https://doi.org/10.1139/o85-102.
(18) Guez, V.; Nair, S.; Chaffotte, A.; Bedouelle, H. The Anticodon-Binding Domain of Tyrosyl-tRNA Synthetase: State of Folding and Origin of the Crystallographic Disorder. *Biochemistry* **2000**, *39* (7), 1739–1747. https://doi.org/10.1021/bi992382v.
(19) Bouvier, M.; Stafford, W. F. Probing the Three-Dimensional Structure of Human Calreticulin. *Biochemistry* **2000**, *39* (48), 14950–14959. https://doi.org/10.1021/bi0019545.
(20) Danielsson, J.; Jarvet, J.; Damberg, P.; Gräslund, A. Translational Diffusion Measured by PFG-NMR on Full Length and Fragments of the Alzheimer Aβ(1–40) Peptide. Determination of Hydrodynamic Radii of Random Coil Peptides of Varying Length. *Magnetic Resonance in Chemistry* **2002**, *40* (13), S89–S97. https://doi.org/10.1002/mrc.1132.
(21) Karlin, D.; Longhi, S.; Receveur, V.; Canard, B. The N-Terminal Domain of the Phosphoprotein of Morbilliviruses Belongs to the Natively Unfolded Class of Proteins. *Virology* **2002**, *296* (2), 251–262. https://doi.org/10.1006/viro.2001.1296.
(22) Longhi, S.; Receveur-Bréchot, V.; Karlin, D.; Johansson, K.; Darbon, H.; Bhella, D.; Yeo, R.; Finet, S.; Canard, B. The C-Terminal Domain of the Measles Virus Nucleoprotein Is Intrinsically Disordered and Folds upon Binding to the C-Terminal Moiety of the Phosphoprotein*. *Journal of Biological Chemistry* **2003**, *278* (20), 18638–18648. https://doi.org/10.1074/jbc.M300518200.
(23) Sánchez-Puig, N.; Veprintsev, D. B.; Fersht, A. R. Human Full-Length Securin Is a Natively Unfolded Protein. *Protein Sci* **2005**, *14* (6), 1410–1418. https://doi.org/10.1110/ps.051368005.
(24) Sánchez-Puig, N.; Veprintsev, D. B.; Fersht, A. R. Binding of Natively Unfolded HIF-1α ODD Domain to P53. *Molecular Cell* **2005**, *17* (1), 11–21. https://doi.org/10.1016/j.molcel.2004.11.019.
(25) Khaymina, S. S.; Kenney, J. M.; Schroeter, M. M.; Chalovich, J. M. Fesselin Is a Natively Unfolded Protein. *J. Proteome Res.* **2007**, *6* (9), 3648–3654. https://doi.org/10.1021/pr070237v.
(26) Zhang, X.; Perugini, M. A.; Yao, S.; Adda, C. G.; Murphy, V. J.; Low, A.; Anders, R. F.; Norton, R. S. Solution Conformation, Backbone Dynamics and Lipid Interactions of the Intrinsically Unstructured Malaria Surface Protein MSP2. *Journal of Molecular Biology* **2008**, *379* (1), 105–121. https://doi.org/10.1016/j.jmb.2008.03.039.
(27) Mittag, T.; Orlicky, S.; Choy, W.-Y.; Tang, X.; Lin, H.; Sicheri, F.; Kay, L. E.; Tyers, M.; Forman-Kay, J. D. Dynamic Equilibrium Engagement of a Polyvalent Ligand with a

Single-Site Receptor. *Proceedings of the National Academy of Sciences* **2008**, *105* (46), 17772–17777. https://doi.org/10.1073/pnas.0809222105.
(28) Paz, A.; Zeev-Ben-Mordehai, T.; Lundqvist, M.; Sherman, E.; Mylonas, E.; Weiner, L.; Haran, G.; Svergun, D. I.; Mulder, F. A. A.; Sussman, J. L.; Silman, I. Biophysical Characterization of the Unstructured Cytoplasmic Domain of the Human Neuronal Adhesion Protein Neuroligin 3. *Biophysical Journal* **2008**, *95* (4), 1928–1944. https://doi.org/10.1529/biophysj.107.126995.
(29) Habchi, J.; Mamelli, L.; Darbon, H.; Longhi, S. Structural Disorder within Henipavirus Nucleoprotein and Phosphoprotein: From Predictions to Experimental Assessment. *PLOS ONE* **2010**, *5* (7), e11684. https://doi.org/10.1371/journal.pone.0011684.
(30) Choi, U. B.; McCann, J. J.; Weninger, K. R.; Bowen, M. E. Beyond the Random Coil: Stochastic Conformational Switching in Intrinsically Disordered Proteins. *Structure* **2011**, *19* (4), 566–576. https://doi.org/10.1016/j.str.2011.01.011.
(31) Perez, R. B.; Tischer, A.; Auton, M.; Whitten, S. T. Alanine and Proline Content Modulate Global Sensitivity to Discrete Perturbations in Disordered Proteins. *Proteins: Structure, Function, and Bioinformatics* **2014**, *82* (12), 3373–3384. https://doi.org/10.1002/prot.24692.
(32) Yarawsky, A. E.; English, L. R.; Whitten, S. T.; Herr, A. B. The Proline/Glycine-Rich Region of the Biofilm Adhesion Protein Aap Forms an Extended Stalk That Resists Compaction. *J Mol Biol* **2017**, *429* (2), 261–279. https://doi.org/10.1016/j.jmb.2016.11.017.
(33) Poznar, M.; Hołubowicz, R.; Wojtas, M.; Gapiński, J.; Banachowicz, E.; Patkowski, A.; Ożyhar, A.; Dobryszycki, P. Structural Properties of the Intrinsically Disordered, Multiple Calcium Ion-Binding Otolith Matrix Macromolecule-64 (OMM-64). *Biochimica et Biophysica Acta (BBA) - Proteins and Proteomics* **2017**, *1865* (11, Part A), 1358–1371. https://doi.org/10.1016/j.bbapap.2017.08.019.
(34) Więch, A.; Rowińska-Żyrek, M.; Wątły, J.; Czarnota, A.; Hołubowicz, R.; Szewczuk, Z.; Ożyhar, A.; Orłowski, M. The Intrinsically Disordered C-Terminal F Domain of the Ecdysteroid Receptor from Aedes Aegypti Exhibits Metal Ion-Binding Ability. *The Journal of Steroid Biochemistry and Molecular Biology* **2019**, *186*, 42–55. https://doi.org/10.1016/j.jsbmb.2018.09.008.
(35) Pesce, F.; Newcombe, E. A.; Seiffert, P.; Tranchant, E. E.; Olsen, J. G.; Grace, C. R.; Kragelund, B. B.; Lindorff-Larsen, K. Assessment of Models for Calculating the Hydrodynamic Radius of Intrinsically Disordered Proteins. *Biophysical Journal* **2023**, *122* (2), 310–321. https://doi.org/10.1016/j.bpj.2022.12.013.
(36) Shaner, N. C.; Campbell, R. E.; Steinbach, P. A.; Giepmans, B. N. G.; Palmer, A. E.; Tsien, R. Y. Improved Monomeric Red, Orange and Yellow Fluorescent Proteins Derived from Discosoma Sp. Red Fluorescent Protein. *Nat Biotechnol* **2004**, *22* (12), 1567–1572. https://doi.org/10.1038/nbt1037.
(37) Cormack, B. P.; Valdivia, R. H.; Falkow, S. FACS-Optimized Mutants of the Green Fluorescent Protein (GFP). *Gene* **1996**, *173* (1), 33–38. https://doi.org/10.1016/0378-1119(95)00685-0.
(38) Zacharias, D. A.; Violin, J. D.; Newton, A. C.; Tsien, R. Y. Partitioning of Lipid-Modified Monomeric GFPs into Membrane Microdomains of Live Cells. *Science* **2002**, *296* (5569), 913–916. https://doi.org/10.1126/science.1068539.
(49) Crameri, A.; Whitehorn, E. A.; Tate, E.; Stemmer, W. P. C. Improved Green Fluorescent Protein by Molecular Evolution Using DNA Shuffling. *Nat Biotechnol* **1996**, *14* (3), 315–319. https://doi.org/10.1038/nbt0396-315.
